# HsbA represses stationary phase biofilm formation in *Pseudomonas putida*

**DOI:** 10.1101/2024.11.25.625174

**Authors:** Marta Pulido-Sánchez, Elisa Montero-Beltrán, Aroa López-Sánchez, Fernando Govantes

**Affiliations:** Centro Andaluz de Biología del Desarrollo, Universidad Pablo de Olavide/Consejo Superior de Investigaciones Científicas/Junta de Andalucía. ES-41013 Sevilla, Spain; Departamento de Biología Molecular e Ingeniería Bioquímica, Universidad Pablo de Olavide. ES-41013 Sevilla, Spain

**Keywords:** Biofilm, c-di-GMP, stationary phase, signal transduction, stress responses, *Pseudomonas*

## Abstract

*Pseudomonas putida* biofilm growth is associated to nutrient-sufficient conditions and biofilm dispersal is induced by nutrient starvation, signaled by the stringent response-associated nucleotide alarmone (p)ppGpp. We have used transcriptomic analysis to show that (p)ppGpp regulates the *hsbAR-hptB* gene cluster, encoding components of a phosphorelay pathway and an anti-σ factor antagonist, and *cfcR*, encoding a response regulator with diguanylate cyclase (DGC) activity. Transcription of *hsbAR-hptB* and *cfcR* is RpoS-dependent and induced by stationary phase and the stringent response. A Δ*hsbA* mutant resumed biofilm formation after dispersal in late stationary phase and displayed increased pellicle formation at the medium-air interphase and Congo Red adsorption. All these phenotypes were traced down to increased c-di-GMP levels in stationary phase, dependent on the activity of CfcR and its cognate sensor kinase, CfcA. HsbA was reversibly phosphorylated by the combined action of HptB and HsbR. HsbA phosphorylation conditioned its interaction with CfcR and CfcA and the subcellular distribution of the three proteins. In spite of this, HsbA retained its ability to prevent biofilm formation regardless of its phosphorylation state. Our results support a model in which HsbA forms a complex with CfcR to inhibit its DGC activity regardless of its phosphorylation state. Upon HsbA dephosphorylation, this complex is recruited to the cell membrane by CfcA to strengthen the inhibitory effect. While this pathway contributes to biofilm dispersal by denying *de novo* c-di-GMP synthesis during nutrient starvation, it may also enable quick restoration of the biofilm phenotype to colonize new sites or during biofilm maturation.

**HIGHLIGHTS:**

- Transcription of *hsbAR-hptB* is activated by the stringent response and RpoS.
- HptB and HsbR control the phosphorylation state of HsbA.
- HsbA prevents biofilm formation in stationary phase.
- HsbA inhibits the DGC activity of CfcR regardless of its phosphorylation state.
- Unphosphorylated HsbA elicits formation of a membrane-bound HsbA-CfcR-CfcA complex.

GRAPHICAL ABSTRACT

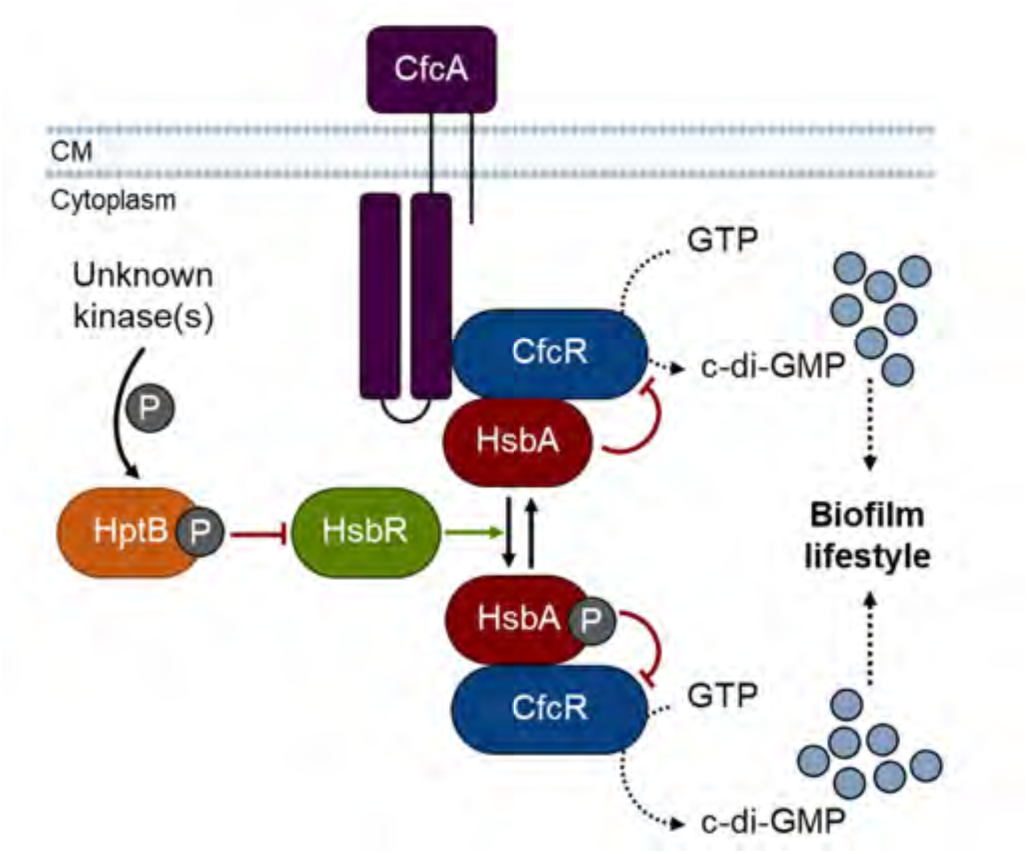

## INTRODUCTION

The life cycles of many bacteria are characterized by the alternation of a single cell-based free-living planktonic stage and a sessile stage during which they develop highly structured surface-associated communities known as biofilms **(Costerton *et al*., 1995)**. Because of the high tolerance to multiple forms of environmental stress associated to the biofilm phenotype, biofilm formation is often regarded as a form of stress response, aimed at protection against harmful conditions, including host defenses and antimicrobial drug challenge. However, biofilm growth may also serve as an efficient means to colonize a favorable niche by providing a nurturing environment and promoting positive interactions between community members **(Davey and O’Toole, 2000; Jefferson *et al*., 2004; McDougald *et al*., 2011)**.

Biofilm formation is a form of coordinated collective behavior regarded as an evolutionary precursor of multicellular development **(Webb *et al*., 2003)**. Bacterial biofilm development proceeds sequentially through stages of attachment, proliferation and maturation, and is terminated by programmed biofilm dispersal to return to the planktonic lifestyle **(O’Toole *et al*., 2000; Monds and O’Toole, 2009; Tolker-Nielsen, 2015)**. Multiple biofilm regulation studies have highlighted the importance of cyclic diguanylate (c-di-GMP)-mediated signaling in establishment, maintenance and termination of the biofilm lifestyle. C-di-GMP is synthesized by diguanylate cyclase (DGC) activities, associated to GGDEF domain proteins, and degraded by phosphodiesterase (PDE) activities, associated to EAL or HD-GYP domain proteins. Bacterial genomes often encode multiple proteins containing these domains, implying that c-di-GMP levels are regulated in a complex fashion. Changes in c-di-GMP concentration are sensed by effector proteins or RNA molecules, which in turn regulate a variety of processes. The biology of c-di-GMP signaling has been reviewed elsewhere **(Boyd and O’Toole, 2012; Römling, 2013; Jenal *et al*., 2017; Hengge, 2021; Homma *et al*., 2022)**.

The Gram-negative saprophytic plant root-associated bacterium *Pseudomonas putida* can form biofilms on a variety of solid substrates, as well as in liquid-air interphases, under nutrient-sufficient conditions. Biofilm formation is signaled by increased c-di-GMP levels that inhibit flagellar gene expression and induce the synthesis of the high molecular weight adhesin LapA, which is required for irreversible surface attachment **(Hinsa *et al*, 2003; Lasa and Penadés, 2006)** and further maintenance of the biofilm structure **(Klausen *et al*., 2006; Martínez-Gil *et al*., 2010)** and extracellular matrix components, such as cellulose **(Navarrete *et al*., 2019)**. Adoption of a biofilm lifestyle provokes profound changes in gene expression, especially in those related to surface properties and motility **(Sauer and Camper, 2001)**. *P. putida* biofilms undergo rapid dispersal in response to carbon or oxygen limitation **(Gjermanssen *et al*., 2005; Hansen *et al*., 2007)**. Starvation-induced biofilm dispersal is effected by proteolytic cleavage of LapA by the periplasmic protease LapG **(Gjermanssen *et al*., 2010)**. Dispersal conditions are signaled by a decrease in the intracellular c-di-GMP concentration caused by activation of the PDE BifA by the stringent response (SR) alarmone (p)ppGpp and the flagellar σ factor FliA **(López-Sánchez *et al*., 2013; Jiménez-Fernández *et al*., 2015; Díaz-Salazar *et al*., 2017)**.

The SR is an adaptive physiological response triggered by different conditions of nutritional or environmental stress and aimed at increasing survival under harsh conditions. The SR is mediated by the synthesis of the hyperphosphorylated guanine nucleotides guanosine 3′,5′-bis(diphosphate) (guanosine tetraphosphate or ppGpp) and guanosine 3′-diphosphate, 5′-triphosphate (guanosine pentaphosphate or pppGpp), collectively known as (p)ppGpp **(Srivatsan and Wang, 2008)**. Increased intracellular (p)ppGpp levels results in major reprogramming of transcription, inhibition of DNA replication, and multiple changes in bacterial metabolism and behavior **(Dalebroux *et al*., 2012; Hauryliuk *et al*., 2015)**. In *P. putida* and other γ- and β-proteobacteria, (p)ppGpp is synthesized by two proteins: RelA, whose activity is induced under amino acid starvation, and SpoT, which responds to other forms of nutritional stress, such as carbon, iron, oxygen or fatty acid limitation **(Gaca *et al*., 2015)**. The general stress response (GSR) is a second adaptive response to multiple environmental stress conditions, including starvation, oxidative conditions, high or low temperature, pH or osmolarity among others **(Bouillet *et al*., 2024)**. In *P. putida* and most γ-proteobacteria, the general stress response is dependent on the alternative σ factor RpoS or σ^S^. RpoS synthesis is heavily regulated in response to different stressors at the levels of transcription, mRNA stability and translation, and its activity is also controlled by proteolysis and by interaction with regulatory factors.

The *hsbA*, *hsbR* and *hptB* genes are highly conserved in the genus *Pseudomonas*, found in the *hsbAR-hptB* arrangement within or in the vicinity of one of the flagellar and chemotaxis gene clusters in most *Pseudomonas* genomes **(Leal-Morales *et al*., 2022)**. HptB, HsbR and HsbA are components of a signal transduction pathway involved in the regulation of flagellar function, biofilm formation and the GSR in *P. aeruginosa* **(Bhuwan *et al*., 2012; Valentini *et al*., 2016; Bouillet *et al*., 2019)**. The single-domain histidine phosphotransferase HptB can be phosphorylated by several hybrid histidine kinases in response to largely unknown signals **(Lin *et al*., 2006; Mern *et al*., 2010; Xu *et al*., 2016; Chen *et al*., 2020)**. HptB can transfer its phosphate group to the response regulator HsbR which, in turn, controls the phosphorylation state of HsbA. Unphosphorylated HsbR is an active kinase that phosphorylates HsbA at the serine-56 residue. Phosphorylation of the HsbR receiver domain by HptB inhibits the kinase and activates the phosphatase activity, leading to HsbA dephosphorylation **(Hsu *et al*., 2008)**. HsbA is a dual function regulator of the bacterial lifestyle. Unphosphorylated HsbA acts as an anti-σ factor antagonist that bind the anti-σ factor FlgM to enable FliA-dependent flagellar gene expression and promote flagellar motility **(Bhuwan *et al*., 2012)**. Phosphorylated HsbA binds HsbR, which is itself an RpoS-specific anti-σ factor, enabling RpoS-dependent activation of the GSR **(Bouillet *et al*., 2019)**. Phosphorylated HsbA also interacts with the membrane-bound DGC HsbD to stimulate c-di-GMP synthesis that leads to upregulation of biofilm formation. Unlike *hsbA*, *hsbR* and *hptB*, *hsbD* is only found in members of the *P. aeruginosa* lineage **(Valentini *et al*., 2016)**.

The *P. putida cfcR* gene encodes a response regulator with an N-terminal receiver domain and an inducible DGC activity **(Matilla *et al*., 2011)**. Expression of *cfcR* is dependent on the GSR σ factor RpoS and activated by Anr and FleQ **(Matilla *et al*., 2011; Ramos-González *et al*., 2016)**, and subjected to posttranscriptional regulation by the RNA-binding proteins RsmA and RsmE **(Huertas-Rosales *et al*., 2017)**. CfcR is activated by phosphorylation by the hybrid histidine kinase CfcA, and the response is stimulated by high salt concentrations **(Ramos-González *et al*., 2016; Tagua *et al*., 2022)**. Deletion of *cfcR* has a modest effect on biofilm formation, but its overexpression causes a pleiotropic phenotype encompassing increased biofilm formation, flocculation and the development of colonies with a crinkly morphology **(Matilla *et al*., 2011; Ramos-González *et al*., 2016)**.

The work presented here demonstrates the function of the HptB-HsbR-HsbA system, in conjunction with the CfcA-CfcR two-component system, in controlling stationary phase biofilm formation in *P. putida*. Our results highlight the use of conserved regulatory elements in different members of the genus *Pseudomonas* to obtain different outcomes that are best suited the physiological demands of each organism.

## MATERIALS AND METHODS

### Bacterial strains and growth conditions

Bacterial strains used in this work are listed in **Table S1**. Liquid cultures of *E. coli* and *P. putida* were routinely grown in lysogeny broth (LB) **(Sambrook and Russell, 2000)** at 37°C and 30°C, respectively, with 180 rpm shaking. Minimal medium was prepared as described **(Mandelbaum *et al*,. 1993)** containing sodium succinate (25 mM) and ammonium chloride (1000 mg l^-1^) as the sole carbon and nitrogen sources. For solid media, American Bacteriological Agar (Condalab) was added to a final concentration of 15 g l^-1^. When needed, antibiotics and other compounds were added at the following concentrations: ampicillin (100 mg l^-1^), carbenicillin (500 mg l^-1^), chloramphenicol (15 mg l^-1^), rifampicin (20 mg l^-1^), gentamycin (10 mg l^-1^), kanamycin (25 mg l^-1^), 3-methylbenzoate (3MB) (3 mM), 5-bromo-4-chloro-3-indoyl-β-D-galactopyranoside (X-gal) (25 mg l^-1^), sodium salicylate (2 mM), Congo Red (40 mg l^-1^) and DL-serine hydroxamate (SHX) (0.8 mM). All reagents were purchased from Sigma-Aldrich.

### Plasmid and strain construction

Strains, plasmids and oligonucleotides used in this work are summarized in **Table S1.** All DNA manipulations were performed following standard protocols **(Sambrook and Russell, 2000)**. Restriction and modification enzymes were used following to the manufacturer’s instructions (New England Biolabs). PCR amplifications were carried out using Q5^®^ High-Fidelity DNA Polymerase (New England Biolabs) for cloning and DreamTaq™ DNA Polymerase (Thermo Fisher) to verify plasmid constructions and chromosomal manipulations. *E. coli* DH5α was used as host strain for cloning procedures. Cloning steps involving PCR were verified by commercial Sanger sequencing (Stab Vida). Plasmid DNA was transferred to *E. coli* strains by transformation **(Inoue *et al*., 1990)**, and to *P. putida* by triparental mating **(Espinosa-Urgel *et al*., 2000)** or electroporation **(Choi *et al*., 2006)**. Site-specific integration of miniTn7-derivatives in *P. putida* strains was performed as previously described **(Choi *et al*., 2005)**, and PCR-verified. Specific details of plasmid and strain construction are provided as **Supplementary Materials and Methods**.

### RNA manipulation for qRT-PCR and RNA-seq

Total RNA from exponential (OD_600_ of 0.4, for qRT-PCR) or stationary phase (OD_600_ of 4.0, for RNA-seq) liquid cultures of *P. putida* was isolated as described **(García-González *et al*., 2005)**. For qRT-PCR experiments, reverse transcription (RT) of 2 µg of total RNA into cDNA was carried out using the High-Capacity cDNA Reverse Transcription kit (Applied Biosystems) following the manufacturer’s instructions. Transcript levels of selected genes were quantified by real-time quantitative PCR (qPCR) using a CFX Connect Real-Time PCR Detection System (Bio-Rad), using appropriate oligonucleotides detailed in **Table S1**. Reactions were prepared using the FastGene^®^ IC Green qPCR Universal kit (Nippon Genetics) following the manufacturer’s instructions.

Samples for RNA-seq experiments were processed at the CABIMER Genomics facility (Centro Andaluz de Biología Molecular y Medicina Regenerativa, Sevilla, Spain) as previously described **(Leal-Morales *et al*., 2022)**.

### SDS-PAGE, Western-blot analysis and co-immunoprecipitation

To evaluate protein expression and stability, SDS-PAGE and Western-blot analyses were performed essentially as described **(Pulido-Sánchez *et al*., 2025)** with some modifications. One-milliliter aliquots from stationary phase cultures (OD_600_ ∼4-5.0) of the wild-type strain bearing the correspondent miniTn*7* derivatives were harvested by centrifugation. Cells were suspended in PAGE loading buffer (80 mM Tris-HCl, pH 6,8; 10% glycerol; 2% SDS; 0.05% bromophenol blue), boiled for 5 minutes and briefly centrifuged. The soluble proteins were separated on 10% SDS-PAGE gels. Blocked nitrocellulose membranes were incubated overnight at 4°C with primary anti-GFP antiserum (SAB4301138, Sigma-Aldrich, 1:5000 dilution), while HRP-conjugated anti-rabbit (31460, Thermo Fischer, 1:10000 dilution) was used as secondary antibody.

To identify interactors of HsbA, HsbA^S56A^ and HsbA^S56D^, we used strains bearing miniTn7 transposons to produce GFP-tagged bait proteins from the P*hsbA* promoter, or solely GFP from the constitutive P*A1/04/03* promoter as negative control. Co-immunoprecipitation assays were subsequently performed using GFP-Trap^®^ agarose beads (ChromoTek). The detailed protocols for protein extraction, immunoprecipitation, protein identification and data analysis are provided as Supplementary Materials and Methods.

### *In vivo* gene expression assays

Overnight LB cultures of strains harbouring transcriptional fusions to *gfp-lacZ* were diluted to an A_600_ of 0.01 in 3 mL of the same medium supplemented with the corresponding antibiotics, and 2 mM or 0.8 mM SHX when required, and incubated at 30 °C with 180 rpm shaking until reaching OD_600_ of 0.5 (exponential phase) or 5.0 (stationary phase). The β-galactosidase activity was determined from sodium dodecyl sulfate- and chloroform-permeabilized cells as previously described **(Miller, 1993)**. For end-point fluorescence measurements, samples were treated and quantified using a Tecan Spark 10M microplate reader as previously described **(Pulido-Sánchez *et al*., 2025)**.

For indirect quantification of c-di-GMP, fluorescence of the strains harbouring the P*cdrA-gfp* transcriptional fusion was monitored along the planktonic curve as previously described **(Leal-Morales *et al*., 2022)**, with the following modifications: (i) overnight LB cultures were 500-fold diluted in 1/10 strength LB medium with appropriate antibiotics and sodium salicylate when needed; (ii) plates were incubated at 30 °C with 180 rpm shaking; and (iii) A_600_ and GFP fluorescence were monitored for 23 h in 15-minute intervals.

### Minimal inhibitory concentration analysis

To determine the minimal concentration of SHX required to inhibit cell growth, overnight LB cultures were 500-fold diluted in minimal medium and 150 µL of the cell suspensions were dispensed into the wells of a Costar^®^ 96-well polystyrene plate (Corning, Ref: 3585). The plate was incubated in a Tecan Spark 10M microplate reader at 30 °C with 180 rpm shaking and OD_600_ was monitored for 23 h in 15-minute intervals. Different concentrations of SHX were added to the wells when cultures reached an OD_600_ of 0.4.

### Pellicle formation assays

To assess the formation of air-liquid interface pellicles, fresh colonies were inoculated in 15 mL glass tubes containing 5 mL of LB medium and incubated at 30 °C with 180 rpm shaking. Sodium salicylate was added to the medium when needed. The presence of glass-attached pellicles was documented after 24 h and 48 h by digital photography.

### Congo Red staining

To evaluate changes in the composition of the extracellular polymeric matrix, Congo Red adsorption assays in plates were performed as described **(Friedman and Kolter, 2004)** with some modifications. Drops of 10 µL of overnight LB cultures were spotted in the surface of T-medium agar plates (1% Bacto^TM^ Tryptone, 1% Bacto^TM^ Agar) supplemented with Congo Red. Sodium salicylate was added to the medium when needed. Plates were incubated at 30 °C for 48 h and colony morphology was documented by digital photography.

### Serial-dilution-based biofilm growth curves

Serial dilution-based biofilm growth on microtiter plates were performed as previously described **(López-Sánchez et al., 2013)**. Plates were incubated for 20 h or 30 h, and sodium salicylate was added to the medium when appropriate. At least three biological replicates were assayed in sextuplicate for each strain.

### Swimming motility assays

Swimming assays were adapted from Parkinson **(Parkinson, 1976)**. Tryptone-agar medium plates containing 0.3% Bacto^TM^ Agar were toothpick-inoculated with fresh-colonies and incubated at 30 °C for 12 h. Sodium salicylate or SHX were added when appropriate. Digital images were taken and swimming halo diameters were measured using GIMP v2.10.10 **(The GIMP Development Team, 2019)** and normalized to the wild-type.

### Confocal microscopy and image analysis

Overnight cultures were 100-fold diluted in LB medium and grown to stationary phase (OD_600_ of 5.0). Cells from 200 µL samples were harvested by centrifugation (5000 *g*, RT, 5 min) and washed three times in PBS to remove LB. For membrane staining, cells were resuspended in 100 µL of PBS containing 5 µg/mL FM™ 4-64 (Molecular Probes). After incubation for 5 min at room temperature in the dark, 2 µL drops were immediately placed on 1% agarose pads made with deionized water to be imaged using an Axio Observer7 confocal microscope (Zeiss) equipped with a CSU-W1 spinning disk module (Yokogawa). Images were taken using a Zeiss α Plan-Apochromat 100x/1.46 Oil DIC M27 objective lens and a Prime 95B CMOS camera (Teledyne Photometrics, pixel size: 0.1099 μm), controlled by the SlideBook 6 software. GFPmut3 fluorescence was excited with the 488 nm laser line and collected with emission filter 525/50 nm, while FM™ 4-64 was excited at 561 nm and collected with a 617/73 filter (exposure time of 1000 ms). Exposure times for GFPmut3-tagged HsbA, HsbA^S56A^, HsbA^S56D^, CfcR and CfcA were 300 ms. For each image, seven Z-sections were collected with a step size of 0.33 μm.

Microscopy images were processed using Fiji software v1.53t **(Schneider *et al*., 2012)** and analyzed using the MicrobeJ v5.13I plugin **(Ducret *et al*., 2016)** following a previously described pipeline **(Pulido-Sánchez *et al*., 2025).**

### Bioinformatics analyses

RNA-seq data treatment and differential expression analyses were performed as previously described **(Leal-Morales *et al*., 2022)**. The raw dataset is available at the Gene Expression Omnibus (GEO) database **(ncbi.nlm.nih.gov/geo)** under accession number GEO: GSE281821.

The intergenic region sequence between *hsbA* (PP_4364) and *fliJ* (PP_4365) from the *P. putida* KT2440 genome was obtained from the *Pseudomonas* Genome Database **(pseudomonas.com)** and manually scanned to detect any σ^S^-dependent promoter consensus sequence **(Peano *et al*., 2015)**.

Protein complex structures were predicted using the AlphaFold3-multimer tool available in alphafoldserver.com **(Abramson *et al*., 2024)**. Protein structures were visualized and analyzed using UCSF ChimeraX v1.11 **(Meng *et al*., 2023)**.

### Statistical analysis

Statistical treatment of the data was performed using GraphPad Prism v8.3.0 software. Unless otherwise stated, results are reported as the average and standard deviation of at least three biological replicates or image datasets, or median and 95% confidence interval for the median in foci fluorescence quantification. Significance (p-values) of the differences among strains/samples was evaluated by means of the two-tailed Student’s t-test for unpaired samples not assuming equal variance. For differential expression analysis of RNA-seq data, p-values were adjusted by the Benjamini-Hochberg method.

## RESULTS

### The *hsbAR-hptB* cluster is regulated by the SR and the GSR

Our previous work showed that *hsbA*, *hsbR* and *hptB* are part of the *hsbAR-hptB-fliKLMNOPQR-flhB* operon of the reference *P. putida* strain KT2440. Transcription of this operon is driven by P*hsbA*, which is modestly activated by FliA, and P*fliK* and P*fliL*, two σ^54^-dependent, FleQ-activated promoters **(Leal Morales *et al*., 2022)**. In a RNA-seq experiment designed to identify genome-wide targets of regulation by the SR-associated alarmone (p)ppGpp, stationary phase *hsbA, hsbR* and *hptB* transcription was increased 17.3-, 13.3- and 3.4-fold, respectively, in the wild-type strain compared to a control Δ*relA*Δ*spoT* strain not producing the alarmone (known as the ppGpp^0^ mutant) **(Fig. 1A)**. None of the downstream genes of the operon or any others within the flagellar gene cluster displayed a similar response **(Figure S1)**, suggesting that stronger transcription from the internal P*fliK* and P*fliL* promoters likely masks this regulation.

**Figure 1.**
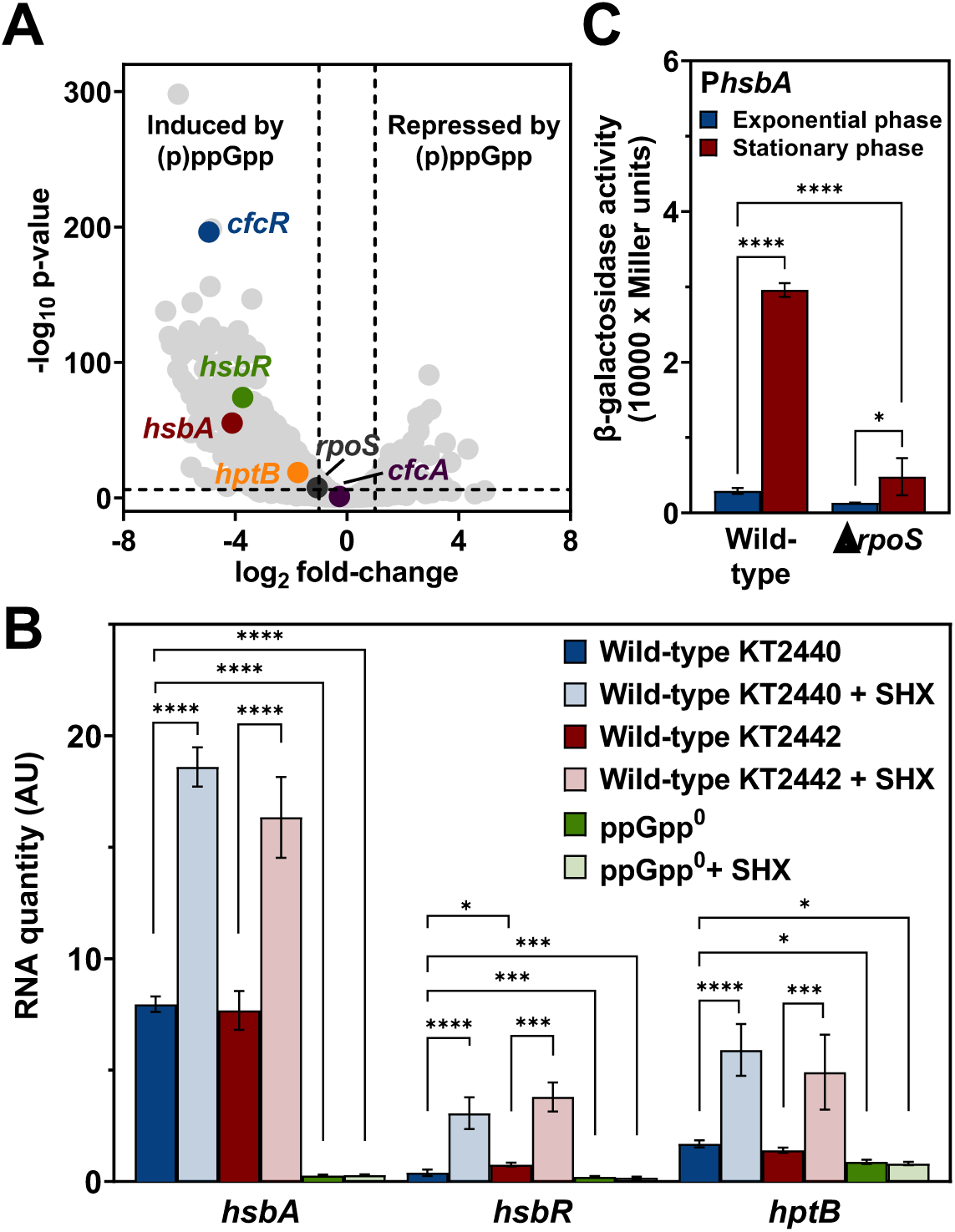
Transcriptional regulation of the *hsbAR-hptB* cluster. **A.** Volcano plot of RNA-seq expression data (GEO:GSE281821) comparing differential gene expression of a ppGpp^0^ mutant relative to the wild-type (KT2440) strain. The horizontal dotted line corresponds to an adjusted p-value of 10^-6^. Vertical dotted lines correspond to a twofold-change expression threshold. Colored points highlight genes assessed in this work. **B.** qRT-PCR quantification of *hsbA, hsbR* and *hptB* expression in wild-type KT2440, wild-type KT2442 and ppGpp^0^ cells harvested in exponential phase 2 h after the addition of 0.8 mM SHX in minimal medium. **C.** β-galactosidase assays of exponential and stationary phase cultures of wild-type (KT2442) and Δ*rpoS* cells bearing a transcriptional P*hsbA-lacZ* fusion. Columns and error bars represent the averages and standard deviations of at least three biological replicates. Stars designate p-values for the Student’s t-test for unpaired samples not assuming equal variance. ns: non-significant; *:p<0.05; **:p<0.01; ***:p<0.001; ****:p<0.0001.

While the SR is naturally induced in stationary phase upon nutrient exhaustion **(Navarro-Llorens *et al*., 2010)**, it can also be induced experimentally in exponential phase by addition of amino acid analogues such as serine hydroxamate (SHX). SHX binds and interferes with seryl-tRNA synthetases leading to arrest of protein synthesis **(Tosa and Pizer, 1971)**, and SHX addition has been shown to trigger (p)ppGpp accumulation in exponential phase cultures of *P. putida*, resulting in growth inhibition **(Vogeleer and Létisse, 2022).** Indeed, exponential growth of *P. putida* KT2440 in minimal medium slowed down upon addition of SHX in a concentration-dependent fashion and was completely inhibited at 0.8 mM SHX **(Fig. S2)**. We have used qRT-PCR to assess the effect of 0.8 mM SHX addition on the transcript levels of *hsbA*, *hsbR* and *hptB* in the wild-type strain KT2440, its isogenic ppGpp^0^ mutant, and KT2442, a Rif^r^ derivative of KT2440 regularly used in our lab as the reference strain **(Fig. 1B)**. Transcription of *hsbA, hsbR* and *hptB* was increased 2.5-, 7.5- and 3.5-fold, respectively, in SHX-treated KT2440 cells relative to the untreated controls. The levels of the three transcripts were very low in the ppGpp^0^ mutant, and were not increased after the addition of SHX. This indicates that SHX induces the expression of *hsbAR-hptB* by promoting (p)ppGpp accumulation in *P. putida*. The results obtained with KT2442 were equivalent to those observed in KT2440, both in SHX-treated and untreated cells. Accordingly, KT2442 was henceforth used as the reference wild-type strain.

Analysis of the intergenic region upstream from *hsbA* revealed a sequence with high similarity to the binding motif of the RNA polymerase loaded with the alternative σ factor RpoS (σ^S^), with perfect conservation of the discriminating −13C, −9C and −5A positions **(Peano *et al*., 2015) (Fig. S3).** RpoS is the central regulator of the GSR in *Escherichia coli* and other bacteria. In order to test the impact of RpoS on *hsbAR-hptB* transcription, we introduced a plasmid-borne fusion of the P*hsbA* promoter to the dual reporter *gfp*mut3-*lacZ* **(Leal-Morales *et al*., 2022)** into the wild-type KT2442 and its isogenic Δ*rpoS* mutant MRB193 and assessed the β-galactosidase activity derived from the fusion in exponential and stationary phase **(Fig. 1C)**. P*hsbA* expression in the wild-type strain was low in exponential phase and greatly (10-fold) induced in stationary phase. Expression in the Δ*rpoS* mutant was 2- and 6-fold lower in exponential and stationary phase, respectively, although a residual 4-fold stationary phase induction was still observed in this background. Taken together, our results indicate that transcription of the *hsbAR-hptB* gene cluster is subjected to dual regulation by the SR and the GSR.

### HsbA is a negative regulator of biofilm formation

Because HsbA is a positive regulator of biofilm formation in *P. aeruginosa* **(Bordi *et al*., 2010; Valentini *et al*., 2016; Bouillet *et al*., 2019)**, we set out to test the impact of HsbA on biofilm formation in *P. putida*. To this end, we constructed the KT2442 Δ*hsbA* mutant derivative MRB171. Deletion of *hsbA* did not cause transcriptional polarity on the downstream genes *hsbR* and *hptB*, as evidenced by qRT-PCR **(Fig. S4)**. Next, we measured submerged biofilm growth, medium-air interphase biofilm (pellicle) formation and colony morphology on agar plates containing Congo Red, an indicator of extracellular polymeric matrix (EPM) production **(Jones and Wozniak, 2017)**, in the wild-type and its Δ*hsbA* mutant derivative MRB171 **(Fig. 2)**. Submerged biofilm growth was assessed by means of dilution series-based growth curves in microtiter plates **(López-Sánchez *et al*., 2013) (Fig. 2A)**. In these conditions, the wild-type strain formed a biofilm during exponential growth and underwent dispersal during stationary phase, as previously described **(López-Sánchez *et al*., 2013)**. The Δ*hsbA* mutant also formed a biofilm during exponential phase and initiated dispersal in early stationary phase with dynamics similar to that of the wild-type. However, dispersal was interrupted and biofilm formation was resumed in late stationary phase. Extended incubation, from 20 to 30 hours, resulted in further biofilm growth of the Δ*hsbA* mutant with no sign of dispersal, while the wild-type strain did not resume biofilm growth. In the medium-air interphase biofilm formation assays, the wild-type strain formed a thin pellicle after 24 h incubation in rich medium **(Fig. 2B)** that was no longer present at 48h. In contrast, the Δ*hsbA* mutant exhibited a thick pellicle that was still evident at 48 h. Finally, the wild-type strain formed smooth colorless colonies on Congo Red agar **(Fig. 2C)**, while colonies of the Δ*hsbA* mutant were stained red with no visible change in morphology. Taken together, these results strongly suggest that under our experimental growth conditions HsbA is a negative regulator that prevents the formation of a submerged biofilm in stationary phase, the production of a robust, dispersal-resistant medium-air biofilm and the synthesis of Congo Red-stainable EPM components during growth on agar plates.

**Figure 2.**
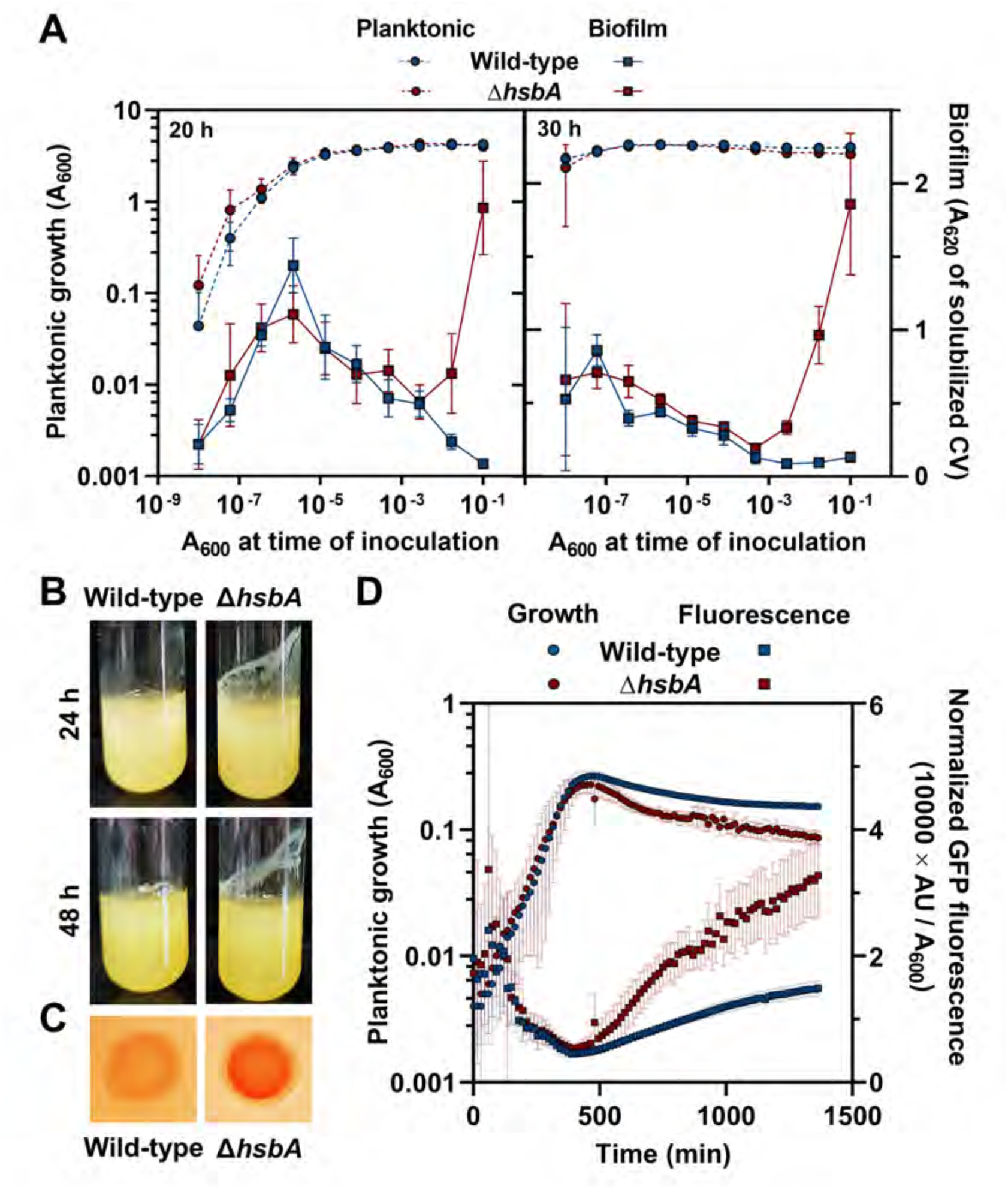
Biofilm-related phenotypes of the Δ*hsbA* mutant. Phenotypic assays of the wild-type and Δ*hsbA* strains. **A.** Dilution series-based growth curves after 20 h (left) or 30 h (right) incubation. Planktonic (left axis, circles) and biofilm growth (right axis, squares) are plotted against the initial A_600_ of each dilution. CV: crystal violet. Points and error bars represent averages and standard deviations of six technical replicates. A representative assay out of at least three biological replicates is shown. **B.** Pellicle formation in glass tubes of LB-cultures after 24 h (top) or 48 h (bottom) incubation. **C.** Congo Red adsorption by colonies after 48 h incubation. **D.** Time course of the fluorescence of the aforementioned strains bearing the c-di-GMP biosensor plasmid pCdrA::*gfp*(ASV)^C^ grown in 1/10 strength LB. Planktonic (left axis, circles) and normalized GFP fluorescence counts (right axis, squares) are plotted against time. Points and error bars represent averages and standard deviations of at least three biological replicates.

As biofilm formation and Congo Red adsorption are generally associated with high c-di-GMP concentrations in the cell **(Jones and Wozniak, 2017)**, we next assessed the c-di-GMP accumulation dynamics in the wild-type and the Δ*hsbA* strains. The pCdrA::*gfp* plasmids are a family of fluorescent c-di-GMP reporters based on the FleQ-repressed and c-di-GMP-inducible P*cdrA* promoter from *P. aeruginosa*. In a FleQ^+^ background, fluorescence levels recapitulate the intracellular c-di-GMP concentration **(Rybtke et al., 2012)**. These plasmids have been widely used in the genus *Pseudomonas* as a convenient alternative to direct c-di-GMP measurements (**López-Farfán *et al*., 2019; Barrientos-Moreno *et al*., 2022; Xiao *et al*., 2022; Scribani-Rossi *et al*., 2023)**, To avoid excessive fluorescence accumulation due to the high stability of GFP, we used plasmid pCdrA::*gfp*(ASV)^C^, in which P*cdrA* drives the synthesis of the unstable GFP(ASV) variant **(Andersen *et al*., 1998)**. The normalized fluorescence levels in the pCdrA::*gfp*(ASV)^C^-bearing wild-type strain decreased during exponential growth, likely due to dilution derived from growth counteracting fluorescence accumulation **(Fig. 2D)**, and increased as growth stopped upon entry in stationary phase. The behavior of the Δ*hsbA* mutant was similar, but the fluorescence signal increased at a higher rate in stationary phase, suggesting that Δ*hsbA* mutant cells bear greater intracellular c-di-GMP concentrations than wild-type cells at this stage. These results strongly suggest that HsbA inhibits biofilm formation in stationary phase by preventing c-di-GMP accumulation.

### HsbA inhibits c-di-GMP synthesis by diguanylate cyclase CfcR

In *P. aeruginosa*, HsbA stimulates c-di-GMP synthesis by interacting with the DGC HsbD **(Valentini *et al*., 2016)**. In contrast, our results suggest that *P. putida* HsbA prevents c-di-GMP synthesis or promotes its degradation, and an HsbD orthologs is not present **(Valentini *et al*., 2016)**. To identify possible targets for HsbA action in *P. putida*, we conducted *in vivo* co-immunoprecipitation (co-IP) assays. To this end, we constructed a miniTn*7* derivative expressing a HsbA-GFP fusion from P*hsbA* and the *hsbA* translational start region. Western blot of the wild-type strain bearing the P*hsbA-hsbA-gfp* mini-Tn*7* derivative revealed a band reactive with GFP antiserum compatible with the expected HsbA-GFP molecular weight (38 kDa), suggesting that the fusion protein is produced and stable **(Fig. S5)**. HsbA-GFP production prevented biofilm regrowth to rescue the wild-type phenotype when produced in the Δ*hsbA* mutant during stationary phase **(Fig. S6)**. Next, cells from stationary phase cultures of the wild-type strain bearing the P*hsbA-hsbA-gfp* transposon were disrupted and the solubilized cell contents were incubated with anti-GFP affinity beads to capture HsbA-GFP and its interaction partners. Co-immunoprecipitated proteins were subsequently analyzed by mass spectrometry. Among the identified proteins, the strongest enrichment scores corresponded to the response regulator HsbR and the diguanylate cyclase CfcR **(Fig. S7 and Supplementary Data SD1)**. CfcR synthesis was previously shown to be dependent on RpoS **(Matilla *et al*., 2011)**. Interestingly, our RNA-seq results show that *cfcR* is also downregulated in the absence of (p)ppGpp **(Fig. 1A)**, and qRT-PCR assays in the presence of SHX indicate that *cfcR* is a target for positive regulation by the SR **(Fig. S8)**. The DGC activity of CfcR was previously shown to be dependent on its phosphorylation by the membrane-bound hybrid histidine kinase CfcA **(Ramos-González *et al*., 2016; Tagua *et al*., 2022)**. Although *cfcA* is transcribed preferentially in stationary phase **(Tagua *et al*., 2022)**, our results show that its expression is not regulated by the SR **(Figs. 1A and S8)**.

In order to determine whether the increase in stationary phase c-di-GMP levels in the absence of HsbA is caused by the DGC activity of CfcR, we constructed KT2442-derived Δ*cfcR* (MRB229), Δ*cfcA* (MRB230), Δ*hsbA*Δ*cfcR* (MRB231) and Δ*hsbA*Δ*cfcA* (MRB232) deletion mutants and characterized their biofilm-related phenotypes as above **(Fig. 3)**. Deletion of *cfcR* or *cfcA* had no impact on submerged biofilm growth, pellicle formation or Congo Red adsorption **(Figs. 3A, B and C)**. Likewise, the c-di-GMP accumulation dynamics in the Δ*cfcR* and Δ*cfcA* mutant was indistinguishable from that of the wild-type strain **(Fig. 3D)**, suggesting that CfcR is not active in these conditions. However, deletion of *cfcR* or *cfcA* suppressed all the identified biofilm-related phenotypes of the Δ*hsbA* mutant, to rescue the wild-type phenotypes, i.e., the Δ*hsbA*Δ*cfcR* and Δ*hsbA*Δ*cfcA* mutants (i) did not resume biofilm growth after dispersal during stationary phase **(Fig. 3A)**; (ii) did not produce a persistent pellicle after 48 hour incubation **(Fig. 3B)**; (iii) failed to adsorb Congo Red on the indicator plates **(Fig. 3C)**; and (iv) displayed similar c-di-GMP levels to the wild-type strain all along the growth curve **(Fig. 3D)**. Based on the epistasis of *cfcR* and *cfcA* over *hsbA*, along with the evidence suggesting physical interaction between HsbA and CfcR **(Fig. S7)** and the dependency of CfcR DGC activity on CfcA-dependent phosphorylation **(Ramos-González *et al*., 2016)**, we propose that HsbA prevents biofilm formation by directly inhibiting the DGC activity of CfcR or its activation by CfcA.

**Figure 3.**
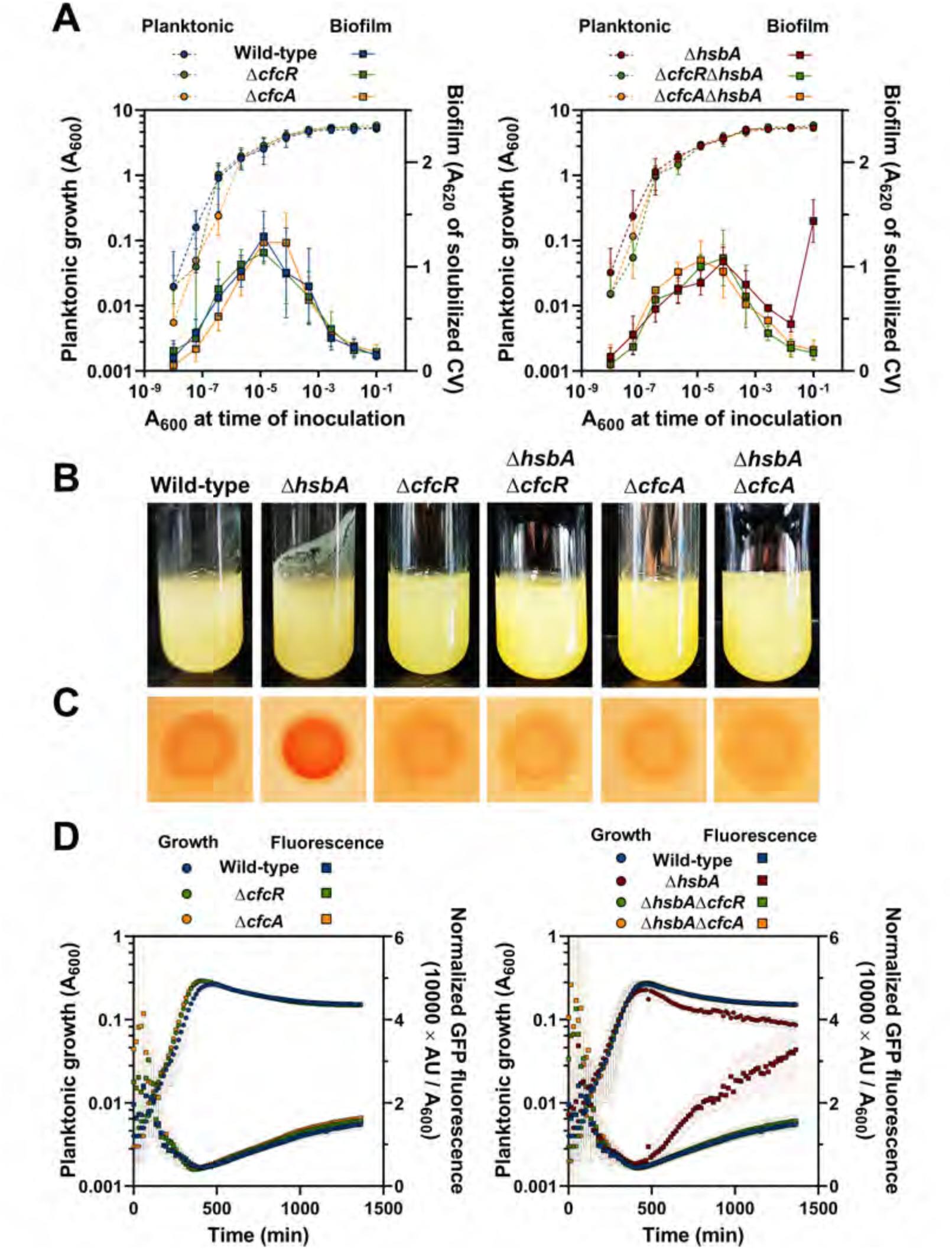
Biofilm-related phenotypes of the Δ*cfcR* and Δ*cfcA* mutants. Phenotypic assays of the Δ*cfcR* and Δ*cfcA* mutations in wild-type and Δ*hsbA* backgrounds. **A.** Dilution series-based growth curves after 20 h incubation of single Δ*cfcR* and Δ*cfcA* mutants *vs.* wild-type strain (left), or double Δ*hsbA*Δ*cfcR* and Δ*hsbA*Δ*cfcA* mutants *vs.* the Δ*hsbA* mutant (right). Planktonic (left axis, circles) and biofilm growth (right axis, squares) are plotted against the initial A_600_ of each dilution. CV: crystal violet. Points and error bars represent averages and standard deviations of six technical replicates. A representative assay out of at least three biological replicates is shown. **B.** Pellicle formation of the aformentioned strains in glass tubes of LB-cultures after 48 h incubation. **C.** Congo Red adsorption by colonies of the aformentioned strains after 48 h incubation. **D.** Time course of the fluorescence of single Δ*cfcR* and Δ*cfcA* mutants *vs.* the wild-type strain (left), or double Δ*hsbA*Δ*cfcRR* and Δ*hsbA*Δ*cfcA* mutants *vs.* the Δ*hsbA* mutant (right), bearing the c-di-GMP biosensor plasmid pCdrA::*gfp*(ASV)^C^ grown in 1/10 strength LB. Planktonic (left axis, circles) and normalized GFP fluorescence counts (right axis, squares) are plotted against time. Points and error bars represent averages and standard deviations of at least three biological replicates.

### CfcA recruits CfcR and HsbA to the cell envelope

CfcA is described as a membrane-associated hybrid histidine kinase with an amino terminal periplasmic CHASE sensor domain flanked by two transmembrane helices, a cytoplasmic GAF sensor domain followed by a histidine autokinase/phosphotransferase domain, and three receiver domains in its carboxyl region **(Tagua *et al*., 2022)**. Because CfcA is located at the cell membrane, we speculated that its interaction with CfcR and/or CfcR interaction with HsbA may condition the cellular location of the three proteins. To test this hypothesis, we constructed miniTn*7* derivatives producing a CfcR-GFP and a CfcA-GFP fusion protein from their natural P*cfcR* and P*cfcA* promoters and translation initiation regions. As above, production and stability of the fusion proteins was verified by Western blot of the wild-type strain bearing the corresponding mini-Tn*7* derivatives, and bands reactive with GFP antiserum compatible with the expected CfcR-GFP and CfcA-GFP molecular weights (106 and 156 kDa, respectively) were observed **(Fig. S5)**. The CfcR-GFP and CfcA-GFP fusions partially restored the pellicle formation phenotype of the Δ*hsbA* strain when expressed from the Δ*hsbA*Δ*cfcR* and Δ*hsbA*Δ*cfcA* mutant chromosomes **(Fig. S9)**, indicating that they retain at least part of the function of the native CfcR and CfcA proteins. Next, we used confocal microscopy to assess the intracellular location of CfcA-GFP, CfcR-GFP and HsbA-GFP in the wild-type, Δ*hsbA*, Δ*cfcR* and Δ*cfcA* strains. A strong fluorescent signal was detected for all proteins in stationary phase cells **(Fig. 4A)**. In these conditions, fluorescence from the CfcA-GFP fusion was diffusely distributed in the wild-type cells. The signal intensity was generally higher in the cell periphery, where 1-2 irregular foci were found in ∼65% of the cells (**Figs. 4A and S10A)**. A dual distribution pattern was also observed with the wild-type strain expressing CfcR-GFP and HsbA-GFP, but peripheral foci were only found in ∼14 and ∼9% of the cells, respectively **(Figs. 4A, S10B and C)**. The distribution of CfcA-GFP was not greatly altered by deletion of *cfcR* or *hsbA*, although the frequency of cells not bearing CfcA-GFP foci was somewhat higher in both backgrounds (**Figs. 4A and S10A)**. In contrast, fluorescent CfcR-GFP foci were not observed in Δ*cfcA* cells and their frequency was decreased 6-fold in the Δ*hsbA* background (**Figs. 4A and S10B)**, while HsbA-GFP foci were absent in the Δ*cfcA* and Δ*cfcR* cells (**Figs. 4A and S10C)**. These results suggest that, under our experimental conditions, CfcA, CfcR and HsbA are partitioned between the cytoplasm and cell envelope-associated complexes. In these conditions, CfcA appears to be essential for the association of CfcR and HsbA with the cell envelope, while CfcR and HsbA have a modest stimulatory effect on the ability of CfcA to nucleate at the cell surface. In addition, CfcR and HsbA are mutually required for their nucleation, suggesting that they are recruited as a CfcR-HsbA complex.

**Figure 4.**
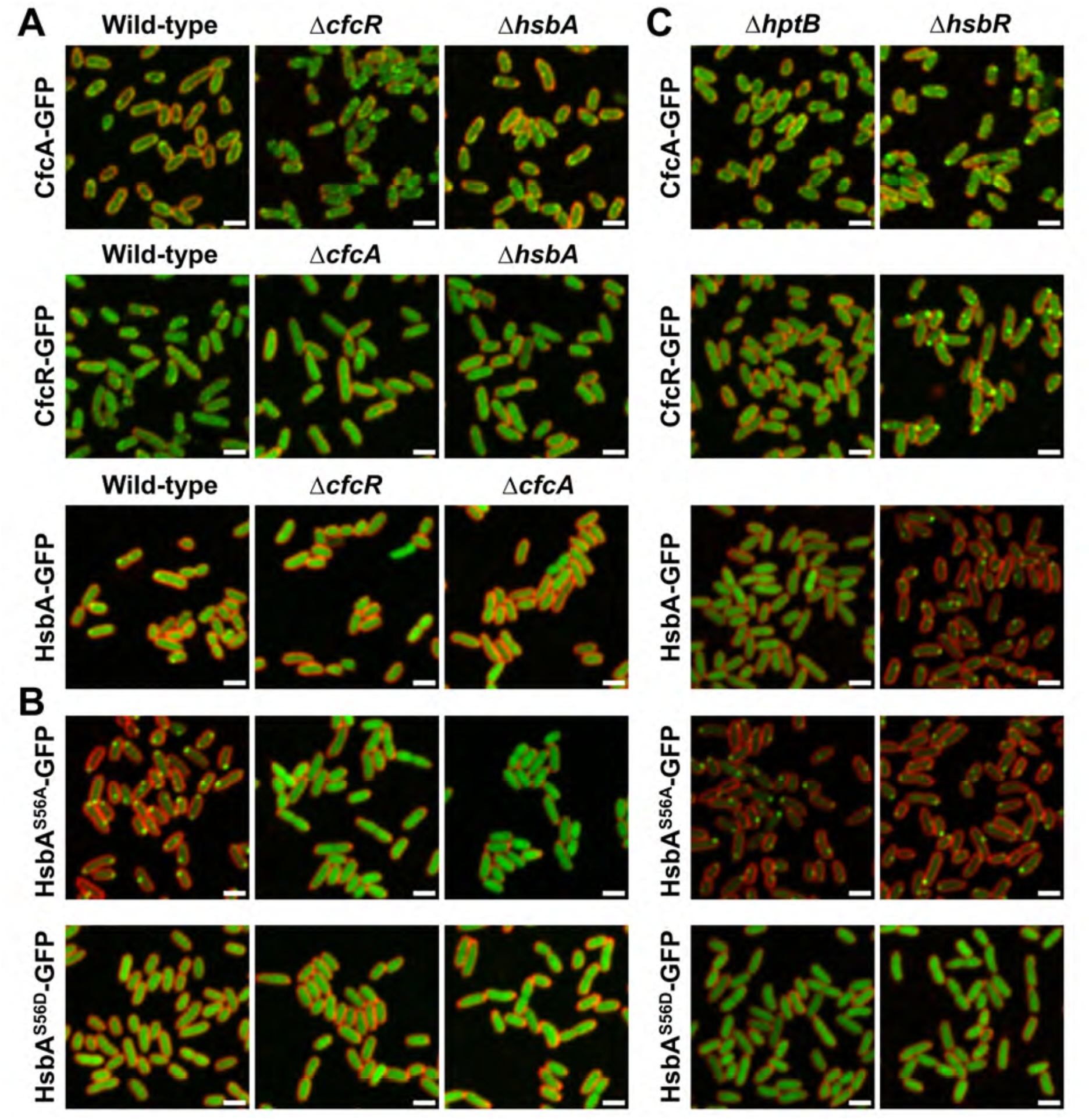
Intracellular localization of CfcA, CfcR and HsbA. Confocal microscopy images of stationary phase wild-type, Δ*cfcR,* Δ*cfcA* or Δ*hsbA* cells bearing the P*cfcA-cfcA-gfp*, P*cfcR-cfcR-gfp* **(A),** or P*hsbA-hsbA-gfp,* P*hsbA-hsbA*^S56A^*-gfp* or P*hsbA-hsbA*^S56D^*-gfp* **(B)** transposons (green). **C.** Confocal microscopy images of stationary phase Δ*hptB* and Δ*hsbR* cells bearing the P*cfcA-cfcA-gfp,* P*cfcR-cfcR-gfp,* P*hsbA-hsbA-gfp,* P*hsbA-hsbA*^S56A^*-gfp* or P*hsbA-hsbA*^S56D^*-gfp* transposons (green). FM^TM^ 4-64 was used as membrane stain (red). Images are shown as the maxima projections of seven Z-sections of the green channel, merged with the red channel showing the cell contour at the confocal plane. Scale bar: 2 µm.

### The intracellular distribution of HsbA, CfcR and CfcA is regulated by HsbA phosphorylation

In *P. aeruginosa*, HsbA activity is regulated by phosphorylation of the conserved serine-56 residue **(Valentini *et al*., 2016)**. Since HsbA displays a dual distribution pattern with most of the protein dispersed in the cytoplasm and a fraction located at discrete positions of the cell envelope, we questioned whether the surface association-dissociation dynamics of HsbA is modulated by its phosphorylation state. To assess this issue, we constructed two more miniTn*7* derivatives producing translational HsbA-GFP fusions in which serine-56 was replaced by alanine (HsbA^S56A^), which prevents phosphorylation at this position, or aspartic acid (HsbA^S56D^), previously shown to mimic the phosphorylated state of its *P. aeruginosa* counterpart **(Valentini *et al*., 2016)**. Production of the fusion proteins was corroborated by Western blot using anti-GFP antiserum **(Fig. S5).** Next, we determined the intracellular distribution of the mutant fusion proteins by confocal microscopy of stationary phase wild-type and mutant cells bearing the corresponding transposons **(Figs. 4B,S10D and E)**. The non-phosphorylatable HsbA^S56A^-GFP fusion was not diffusely distributed in the cytoplasm; rather, 1-3 discrete foci were found at the cell periphery in >98% of the wild-type cells. In contrast, the phosphomimetic HsbA^S56D^-GFP fusion displayed a diffuse distribution and no foci were observed, suggesting that envelope association is specific to the non-phosphorylated form of HsbA. A diffuse distribution was obtained with both HsbA variants in the Δ*cfcA* and Δ*cfcR* backgrounds, indicating that association of the non-phosphorylatable protein to the cell surface is still dependent of the presence of CfcA and CfcR. To test whether the two forms of HsbA display a differential interaction pattern, we performed co-IP experiments as above using extracts from stationary phase wild-type cells producing HsbA^S56A^-GFP or HsbA^S56D^-GFP **(Fig. S11 and Supplementary Data SD1)**. CfcA was specifically co-precipitated by the non-phosphorylatable HsbA^S56A^-GFP, while CfcR was co-precipitated with both variants, albeit with less efficiency by HsbA^S56D^-GFP. Taken together, our results suggest that non-phosphorylated HsbA is specifically recruited to the cell envelope by membrane-bound CfcA only in the presence of CfcR.

In *P. aeruginosa*, the phosphorylation state of HsbA is modulated by the HptB-HsbR phosphorelay pathway **(Hsu et al., 2008; Lin et al., 2006)**. Accordingly, *hsbR* deletion results in constitutively unphosphorylated HsbA, while *hptB* deletion leads to constitutive phosphorylation of HsbA **(Bhuwan *et al*., 2012; Valentini *et al*., 2016)**. HptB, HsbR and several of the hybrid kinases are conserved in *P. putida*, and our results above show that HsbR co-precipitates with HsbA-GFP and HsbA^S56A^-GFP **(Figs. S7 and S11A)**. To test the impact of HptB and HsbR on the interactions between HsbA, CfcR and CfcA, we constructed KT2442-derived in-frame *hptB* and *hsbR* deletion mutants MRB178 and MRB179, and determined the intracellular location of the HsbA-GFP, CfcA-GFP, CfcR-GFP, HsbA^S56A^-GFP and HsbA^S56D^-GFP fusions in these backgrounds **(Figs. 4C and S10)**. The wild-type HsbA-GFP protein fusion was diffusely distributed in the Δ*hptB* mutant cells, but appeared exclusively as 1-3 envelope-associated foci in 99% of the Δ*hsbR* cells, as shown above for the phosphomimetic and non-phosphorylatable HsbA^S56D^ and HsbA^S56A^ variants, respectively. In contrast, localization of HsbA^S56A^-GFP and HsbA^S56D^-GFP was unaffected by the Δ*hsbR* or Δ*hptB* mutations, consistent with the inability of these variants to undergo phosphorylation or dephosphorylation. The CfcR-GFP fusion displayed a similar behavior to HsbA-GFP: a diffuse cytoplasmic distribution was observed in ∼96% Δ*hptB* mutant cells, but it appeared exclusively as envelope-associated foci in ∼91% Δ*hsbR* mutant cells. Finally, distribution of the CfcA-GFP fusion was modestly altered by the Δ*hptB* and Δ*hsbR* mutations, displaying generalized diffuse cytoplasmic distribution with somewhat lower and somewhat higher frequencies of cells bearing peripheral foci, respectively, in these backgrounds. In addition, the CfcA-GFP foci were more defined and displayed higher fluorescence intensity in the absence of HsbR **(Figs. 4C, S10A and S12)**, suggesting that unphosphorylated HsbA may stimulate CfcA nucleation at the cell membrane. Taken together, our results indicate that HptB and HsbR regulate HsbA intracellular distribution by controlling its phosphorylation/dephosphorylation balance, with the phosphorylated form displaying a diffuse cytoplasmic location and the unphosphorylated form being a part of a CfcA- and CfcR-bound membrane-associated complex. In addition, unphosphorylated HsbA is also essential to recruitment of CfcR and stimulates CfcA localization to form a membrane-bound tripartite CfcA-HsbA-CfcR complex.

### Regulation of biofilm formation is not dependent on the HsbA phosphorylation state

To test the impact of HsbA phosphorylation on the regulation of biofilm formation, we assessed the biofilm-related phenotypes observed above in *P. putida* strains lacking HptB or HsbR, which are expected to bear only the phosphorylated or unphosphorylated form of HsbA, respectively **(Fig. 5)**. The *ΔhptB* and Δ*hsbR* strains displayed phenotypes indistinguishable from the wild-type in all assays, i.e., (i) a submerged biofilm was formed in exponential phase, dispersed in stationary phase and not resumed in late stationary phase (**Fig. 5A)**, (i) a thick pellicle was not observed at the medium-air interphase **(Fig. 5B)**, (iii) the *ΔhptB* and Δ*hsbR* colonies did not significantly adsorb the Congo Red dye **(Fig. 5C)**, and (iv) expression levels of the c-di-GMP reporter pCdrA::*gfp*(ASV)^C^ were not increased in stationary phase above those observed in the wild-type **(Fig. 5D)**. These results are in sharp contrast with the differential behavior displayed by both mutants in the localization of HsbA, CfcR and CfcA, and suggest that HsbA phosphorylation has no impact on its ability to regulate biofilm formation. To test this hypothesis further, we constructed miniTn*7* derivatives expressing wild-type HsbA and the HsbA^S56A^ and HsbA^S56D^ variants from the salicylate-inducible P*sal* promoter and tested the ability of these constructs to complement the biofilm formation phenotypes of the Δ*hsbA* mutant. In the absence of salicylate, the presence of HsbA or its mutant variants failed to modify drastically the behavior of the Δ*hsbA* mutant **(Fig. 6)**. In contrast, complementation to different extents was observed in the presence of salicylate. Firstly, biofilm formation was not resumed in stationary phase in the presence of the wild-type HsbA or the HsbA^S56A^ and HsbA^S56D^ proteins **(Fig. 6A)**. Secondly, a thin pellicle, indicative of partial complementation of the Δ*hsbA* mutant phenotype, was observed with the three HsbA variants **(Fig. 6B)**. Thirdly, Congo Red adsorption was abolished in the presence of the three HsbA variants, leading to a colony phenotype similar to the wild-type in all cases **(Fig. 6C)**. Finally, expression of the pCdrA::*gfp*(ASV)^C^ reporter in the presence of the HsbA and HsbA^S56A^ proteins was similar to that of the wild-type strain, but intermediate expression levels between the wild-type and the Δ*hsbA* mutant when HsbA^S56D^ was expressed **(Fig. 6D)**. Taken together, these results suggest that HsbA is competent to repress stationary phase biofilm formation in both its unphosphorylated and phosphorylated form. However, the differences in subcellular localization of phosphorylated and unphosphorylated HsbA and their impact on CfcR and CfcA location, along with minor differences in the extent of complementation of the Δ*hsbA* mutant phenotypes suggest that the two forms may operate through somewhat different mechanisms.

**Figure 5.**
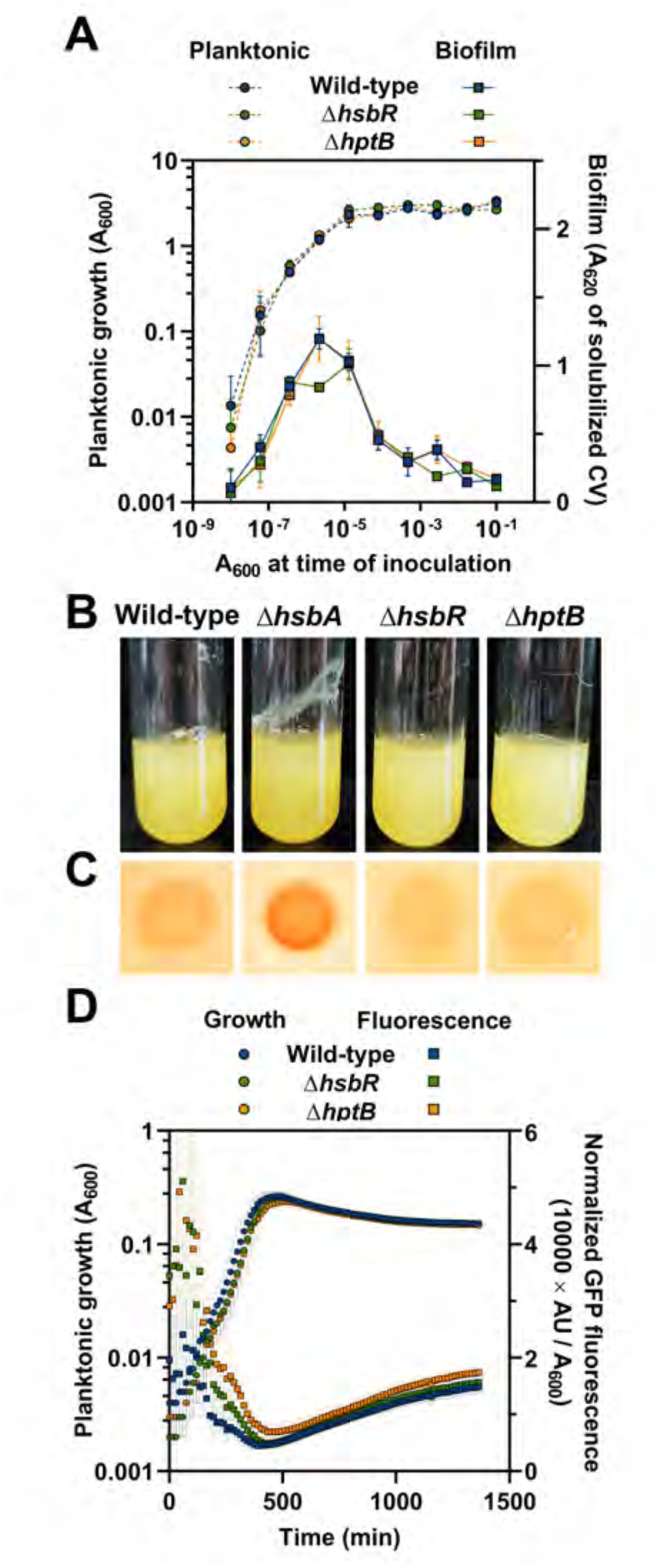
Biofilm-related phenotypes of the Δ*hsbR* and Δ*hptB* mutants. Phenotypic assays of the wild-type, Δ*hsbR* and Δ*hptB* strains. **A.** Dilution series-based growth curves after 20 h incubation. Planktonic (left axis, circles) and biofilm growth (right axis, squares) are plotted against the initial A_600_ of each dilution. CV: crystal violet. Points and error bars represent averages and standard deviations of six technical replicates. A representative assay out of at least three biological replicates is shown. **B.** Pellicle formation in glass tubes of LB-cultures after 48 h incubation. **C.** Congo Red adsorption by colonies after 48 h incubation. **D.** Time course of the fluorescence of the aforementioned strains bearing the c-di-GMP biosensor plasmid pCdrA::*gfp*(ASV)^C^ grown in 1/10 strength LB. Planktonic (left axis, circles) and normalized GFP fluorescence counts (right axis, squares) are plotted against time. Points and error bars represent averages and standard deviations of at least three biological replicates.

**Figure 6.**
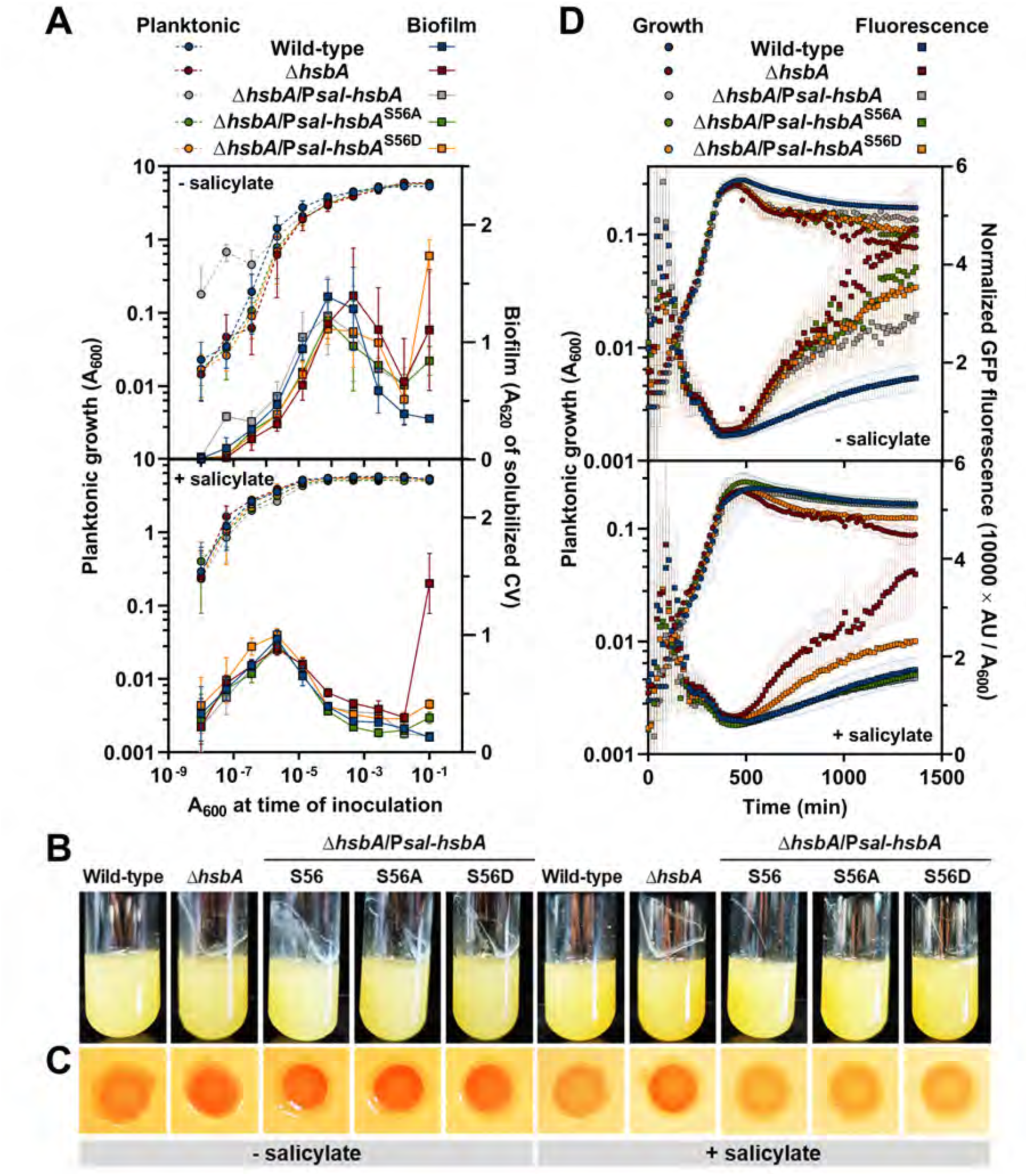
Biofilm-related phenotypes of the non-phosphorylatable and phosphomimetic HsbA variants. Phenotypic assays of the wild-type, Δ*hsbA* mutant, and the Δ*hsbA* mutant expressing wild-type HsbA, HsbA^S56A^ or HsbA^S56D^ from the P*sal* promoter in the absence or in the presence of 2 mM salicylate. **A.** Dilution series-based growth curves after 20 h incubation. Planktonic (left axis, circles) and biofilm growth (right axis, squares) are plotted against the initial A_600_ of each dilution. CV: crystal violet. Points and error bars represent averages and standard deviations of six technical replicates. A representative assay out of at least three biological replicates is shown. **B.** Pellicle formation in glass tubes of LB-cultures after 48 h incubation. **C.** Congo Red adsorption by colonies after 48 h incubation. **D.** Time-course of the fluorescence of the aforementioned strains bearing the c-di-GMP biosensor plasmid pCdrA::*gfp*(ASV)^C^ grown in 1/10 strength LB. Planktonic (left axis, circles) and normalized GFP fluorescence counts (right axis, squares) are plotted against time. Points and error bars represent averages and standard deviations of at least three biological replicates.

### The HptB-HsbR-HsbA system does not regulate flagellar motility in *P. putida*

The HptB-HsbR-HsbA system has also been proposed to regulate flagellar motility in *P. aeruginosa* be means of a partner-switching mechanism. We have tested the impact of *hsbA*, *hsbR* and *hptB* deletions on flagellar swimming in a soft agar-based assay in the presence and in the absence of the SR inducer SHX and found that flagellar function was not significantly impaired in any of the mutant strains **(Figs. 7A and S13)**. Additionally, we tested the Δ*hsbA* mutant complemented with HsbA, HsbA^S56A^ or HsbA^S56D^ produced from the P*hsbA* promoter in the presence or in the absence of SHX, or the three HsbA variants produced from P*sal* in the presence or in the absence of salicylate with similar results **(Figs. 7A and S13)**. These observations indicate that neither elimination of HsbA nor its production at physiological or elevated levels has a significant impact on flagellar function in any of the tested conditions, independently of its phosphorylation state.

**Figure 7.**
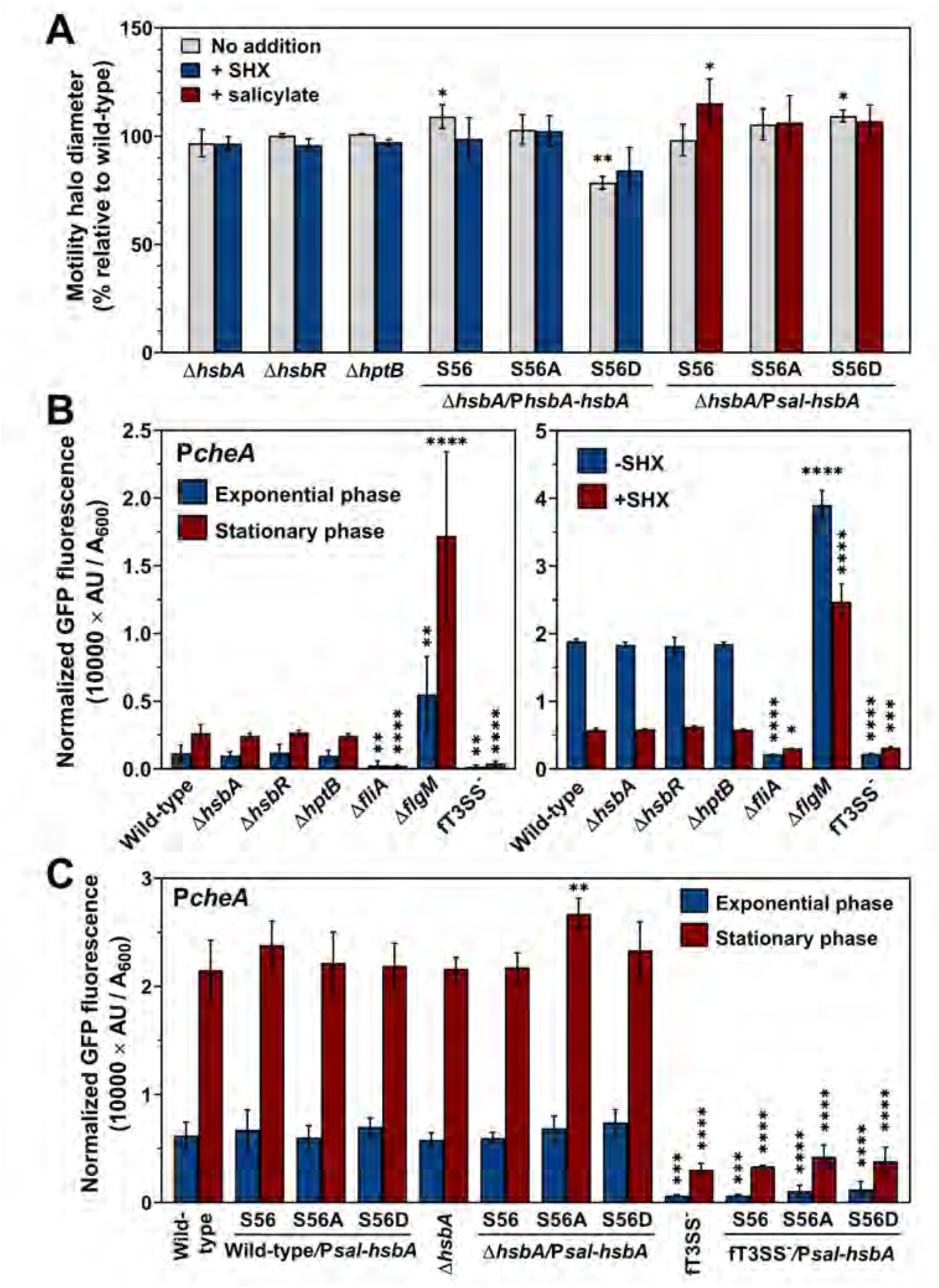
Analysis of flagellar motility and flagellar gene expression. **A.** Relative swimming motility halo diameters of the Δ*hsbA*, Δ*hsbR* and Δ*hptB* mutants, and Δ*hsbA* mutant bearing the P*hsbA-hsbA* (S56), P*hsbA-hsbA*^S56A^ (S56A) or P*hsbA-hsbA*^S56D^ (S56D) transposons in the absence or the presence of 0.8 mM SHX, and the Δ*hsbA* mutant bearing the P*sal-hsbA* (S56), P*sal-hsbA*^S56A^ (S56A) or P*sal-hsbA*^S56D^ (S56D) transposons in the absence or the presence of 2 mM salicylate. Values are relative to the wild-type strain (100%). **B.** Normalized fluorescence from exponential or stationary phase LB cultures (left) or exponential phase cultures in minimal media in the absence of SHX or 2 h after the addition of 0.8 mM SHX (right) of the wild-type, Δ*hsbA*, Δ*hsbR,* Δ*hptB* Δ*fliA*, Δ*flgM* and fT3SS^-^ strains bearing a transcriptional P*cheA-gfp* fusion. **C.** Normalized fluorescence from exponential or stationary phase LB cultures of the wild-type, Δ*hsbA* and fT3SS^-^ strains bearing the P*sal-hsbA,* P*sal-hsbA*^S56A^ or P*sal-hsbA*^S56D^ transposons in the presence of 2 mM salicylate. Columns and error bars represent the averages and standard deviations of at least three biological replicates. Stars designate p-values for the Student’s t-test for unpaired samples not assuming equal variance. ns: non-significant; *:p<0.05; **:p<0.01; ***:p<0.001; ****:p<0.0001.

According to observations in *P. aeruginosa* **(Bhuwan *et al*., 2012)**, we expected an increase in FliA-dependent transcription under conditions that promote HsbA dephosphorylation. To test this possibility, we used a transcriptional fusion of the FliA-dependent P*cheA* promoter to *gfp*mut3 and *lacZ* as a reporter of FliA activity on backgrounds lacking or expressing different levels of HsbA and its variants. Using fluorescence measurements, we determined that P*cheA* transcription was induced in stationary phase, strictly FliA-dependent and upregulated in the absence of FlgM **(Fig. 7B)**. P*cheA* expression levels in a mutant defective in flagellar type III secretion system (fT3SS)-dependent secretion (fT3SS^-^) were low and comparable to those in the Δ*fliA* mutant, consistent with the release of FliA from inactivation by FlgM secretion *via* the flagella **(Pulido-Sánchez *et al*., 2025)**. The fluorescence levels of the P*cheA-gfp-lacZ* fusion in the Δ*hsbA*, Δ*hsbR* and Δ*hptB* mutants were indistinguishable from those in the wild-type strain, and SHX addition decreased P*cheA* expression that was still unaltered by *hsbA*, *hsbR* or *hptB* deletion. We also tested the impact of HsbA, HsbA^S56A^ and HsbA^S56D^ overproduction from the induced P*sal* promoter, but again expression was similar to that in the wild-type strain **(Fig. 7C)**. Finally, we hypothesized that fT3SS-dependent secretion of FlgM might compete with HsbA-dependent FlgM sequestration, effectively masking its impact. This was tested by overproducing HsbA, HsbA^S56A^ and HsbA^S56D^ from P*sal* in the fT3SS^-^ mutant. Again, P*cheA* expression was not altered by production of any of the HsbA variants **(Fig. 7C)**. Taken together, our results indicate that HsbA does not influence FliA-dependent transcription in any of its forms under a variety of conditions, and therefore, it is highly unlikely that HsbA acts as a FlgM antagonist in the regulation of FliA activity in *P. putida*.

## DISCUSSION

The ability to alternate between a planktonic state and the formation of structured surface-associated communities known as biofilms is a major trait of the lifestyles of many bacteria. The Gram-negative soil bacterium *P. putida* is characterized by its ability to develop a biofilm under growth-promoting conditions, while biofilm dispersal is induced by nutrient starvation **(Gjermansen *et al*., 2005)**. In this work we present the interaction of the HptB-HsbR-HsbA signal transduction pathway with the two-component system CfcA-CfcR to contribute to the starvation-induced dispersal response by actively preventing *de novo* biofilm formation under nutritional stress.

The *hsbA*, *hsbR* and *hptB* genes are highly conserved in the genus *Pseudomonas*. They are found in the vast majority of sequenced genomes, invariably linked to gene clusters encoding components of the flagellar and chemotaxis machinery **(Valentini *et al*., 2016; Leal-Morales *et al*., 2022)**. In *P. putida*, *hsbAR-hptB* are transcribed from a promoter region upstream from *hsbA*, designated P*hsbA.* Transcription proceeds into the downstream *fliKLMNOPQR-flhB* flagellar genes, although transcript levels of the distal genes are boosted by two internal promoters upstream from *fliK* and *fliL* **(Leal-Morales *et al*., 2022)**. We show here that *hsbAR-hptB* transcription, but not that of the rest of genes of the operon or any other components of the flagellar cluster, is specifically induced by (p)ppGpp in stationary phase and in response to the SR inducer SHX **(Figs. 1A and C)**. In addition, most transcription from the P*hsbA* promoter region is dependent on the GSR σ factor RpoS and a putative RpoS-dependent promoter motif was readily identified in this region **(Figs. 1B and S3)**. The presence of a motif resembling the consensus RpoS-binding sequence upstream from *hsbA* **(Fig. S3)** suggests that RpoS directly stimulates *hsbAR-hptB* transcription, but we have not experimentally validated this hypothesis yet. We also note that *rpoS* transcription was modestly induced (2.1-fold) in our RNA-seq analysis of (p)ppGpp regulation **(Fig. 1A)**. In addition, (p)ppGpp may also regulate RpoS synthesis and activity posttranscriptionally, as previously shown in *E. coli* **(Battesti *et al*., 2011; Bouillet *et al*., 2024)**. Hence, the effect of (p)ppGpp on P*hsbA* transcription may at least in part be indirect and mediated by RpoS. We previously reported the presence of an additional weak, FliA-dependent promoter in this region **(Leal-Morales *et al*., 2022)** that is likely responsible of the residual activity observed in the absence of RpoS, suggesting that both σ factors contribute to *hsbAR-hptB* transcription. Our previous work showed that the *P. putida* P*bifA* promoter, driving transcription of the c-di-GMP phosphodiesterase responsible for the activation of starvation-induced biofilm dispersal, is also regulated by both FliA and (p)ppGpp **(Díaz-Salazar *et al*., 2017)**. The interaction of nutritional stress regulation with a flagella-associated σ factor on these genes can be envisioned as a means to couple biofilm dispersal with *de novo* flagella synthesis and resumption of the planktonic lifestyle.

HsbR and HsbA display homology to signal transduction proteins that participate in a regulatory mechanism known as the partner switching system (PSS). A PSS is made up of a σ factor, an anti-σ factor with serine kinase activity, a phosphorylatable anti-σ antagonist and a PP2C serine phosphatase. Typically, a PSS anti-σ factor binds its cognate σ factor to prevent it from interacting with RNA polymerase and phosphorylates the anti-σ antagonist, rendering it inactive. In response to a specific physiological or environmental cue, the PP2C phosphatase dephosphorylates the anti-σ antagonist, which can bind and sequester the anti-σ factor, thus releasing the active σ **(Bouillet *et al*., 2018)**. While HsbA displays high similarity with PSS anti-σ factor antagonists **(Bhuwan *et al*., 2012)**, HsbR is a modular protein with significant homology with the *Shewanella oneidensis* anti-σ factor CrsR. Both proteins bear an N-terminal response regulator receiver domain, a central domain with PP2C phosphatase activity and a C-terminal kinase/anti-σ factor domain **(Bouillet *et al*., 2019)**. In *P. aeruginosa*, HsbR controls the phosphorylation state of HsbA. Unphosphorylated HsbR is an active kinase that phosphorylates HsbA at the serine-56 residue. Phosphorylation of the HsbR receiver domain by HptB inhibits the kinase and activates the phosphatase activity, leading to HsbA dephosphorylation **(Valentini *et al*., 2016)**. In this organism, HsbA is a phosphorylation-regulated dual anti-σ factor antagonist: phosphorylated HsbA binds HsbR, which is proposed to act as an anti-σ factor for RpoS **(Bouillet *et al*., 2019)**, while unphosphorylated HsbA interacts with FlgM, the anti-σ factor of FliA **(Bhuwan *et al*., 2012)**. We have extensively tested the possibility that HsbA acts as a functional FlgM antagonist in *P. putida* by assessing flagellar motility phenotypes and expression of the FliA-dependent P*cheA* promoter in a variety of genetic and environmental conditions, including individual deletion of *hsbA*, *hsbR* or *hptB*, ectopic production of phosphomimetic and non-phosphorylatable variants of HsbA, exponential and stationary phase and artificial SR induction with the serine analog SHX. Under none of the conditions tested did we observe swimming behavior or P*cheA* expression levels significantly differing from the wild-type control **(Figs. 7 and S13)**. In addition, co-immunoprecipitation revealed interaction of HsbA with HsbR, CfcR and CfcA, but not FlgM **(Figs. S7 and S11 and Supplementary Data SD1)**. Therefore, we must conclude that the HptB-HsbR-HsbA system is highly unlikely to regulate FliA activity as proposed in *P. aeruginosa*. We have also explored a possible role of HsbA and HsbR in the control of RpoS. Unlike previously reported in *P. aeruginosa* **(Bouillet *et al*., 2019)**, co-IP assays showed an interaction between HsbR and the non-phosphorylatable HsbA^S56A^ derivative, but not with the phosphomimetic HsbA^S56D^ mutant **(Fig. S11)**. On the other hand, analysis of the P*hsbA-lacZ* fusion revealed that *hsbA*, *hsbR* and *hptB* deletions do not affect transcription from the RpoS-dependent P*hsbA* promoter in exponential or stationary phase **(Fig. S14)**. Although this characterization is quite preliminary, our current evidence does not favor the involvement of the HptB-HsbA-HsbR system in the functional regulation of the GSR σ factor RpoS. Whether any of the >20 additional σ factors in *P. putida* is a target for HsbA-dependent partner-switching regulation remains an open question.

Stationary phase and nutrient limitation are associated to biofilm dispersal in *P. putida* **(López-Sánchez *et al*., 2013; Jiménez-Fernández *et al*., 2015)**. We have provided evidence that, in the same conditions, HsbA acts as a negative regulator of biofilm formation. **(Fig. 2)**. HsbA regulation of biofilm formation is dependent on the activity of the CfcA-CfcR two-component system. CfcA is a membrane-bound hybrid histidine kinase and CfcR is a response regulator with DGC activity. CfcR is activated by CfcA-mediated phosphorylation **(Matilla *et al*., 2011; Ramos-González *et al*., 2016; Tagua *et al*., 2022)**. Similarly to HsbA, CfcA and CfcR are synthesized in stationary phase, and *cfcR* transcription is strictly dependent on RpoS **(Matilla *et al*., 2011)** and induced by the SR (**Figs. 1A and S8)**. Our results show that CfcR is co-immunoprecipitated with HsbA **(Fig. S7)**. Deletion of *cfcA* or *cfcR* suppressed all biofilm-related phenotypes of the Δ*hsbA* mutant, but these mutants did not display a phenotype in any of these assays in an HsbA^+^ background, indicating that CfcR is inactive in the presence of HsbA **(Fig. 3)**. While previous reports demonstrated a modest but detectable effect of *cfcR* deletion on biofilm formation **(Matilla *et al*., 2011)**, this apparent contradiction is likely due to different growth conditions and measurement methods. Taken together, our results indicate that HsbA is a *bona fide* component of the starvation-induced biofilm dispersal response of *P. putida*. The activity of HsbA appears to be complementary to that of the PDE BifA: while BifA depletes intracellular c-di-GMP to initiate dispersal **(Jiménez-Fernández *et al*., 2015)**, HsbA actively prevents resumption of biofilm formation by inhibiting *de novo* c-di-GMP synthesis by CfcR.

HsbA and CfcR share a similar dual intracellular location pattern in which a fraction of each protein forms foci at the cell surface, and the rest is diffusely distributed in the cytoplasm. Association with the cell surface was strictly dependent on the presence of CfcA, which itself partitions between the bulk cytoplasm and irregular surface-associated foci. While we have not directly assessed the *in vivo* phosphorylation state of HsbA, a plethora of observations from imaging analysis in wild-type and mutant backgrounds and co-IP analysis suggest that the dynamic intracellular distribution of both proteins is regulated by the phosphorylation state of HsbA. Firstly, deletion of *hptB* or *hsbR* prevent and stimulate the localization of HsbA and CfcR at surface-associated foci, respectively **(Fig. 4C)**. Secondly, the non-phosphorylatable and phosphomimetic HsbA^S56A^ and HsbA^S56D^derivatives phenocopied the HsbA localization in the Δ*hsbR* and the Δ*hptB* mutant, respectively **(Figs. 4 and S10)**. Thirdly, co-IP experiments revealed the interaction of both HsbA derivatives with CfcR, while CfcA was only detected with the non-phosphorylatable HsbA^S56A^ **(Fig. S11)**. Finally, deletion of *hsbR* but not *hptB* caused CfcA to form foci that were more well-defined and displayed higher fluorescence **(Figs. 4C and S12)**. Our results are consistent with a model in which HsbA phosphorylation is controlled by the HptB-HsbR phosphorelay cascade as described in *P. aeruginosa*, where HsbR inactivation prevents HsbA phosphorylation and HptB inactivation results in permanently phosphorylated HsbA **(Hsu *et al*., 2008; Bhuwan *et al*., 2012)**. HsbA and CfcR form a complex that is diffusely located in the cytoplasm when HsbA is in its phosphorylated form, but is recruited by membrane-bound CfcA upon HsbA dephosphorylation **(Fig. 8)**. The HptB-HsbR-HsbA cascade was also shown to influence the intracellular location of its target DGC HsbD in *P. aeruginosa*. HsbD is located at one or both cell poles even in the absence of its N-terminal transmembrane domains and *hptB* deletion boosts polar location, resulting in an increased frequency of cells displaying bipolar HsbD distribution **(Valentini *et al*., 2016)**. Since only the phosphorylated form of HsbA interacts with HsbD in *P. aeruginosa*, this increased polar recruitment is likely a consequence of HsbA hyperphosphorylation in the absence of HptB.

**Figure 8.**
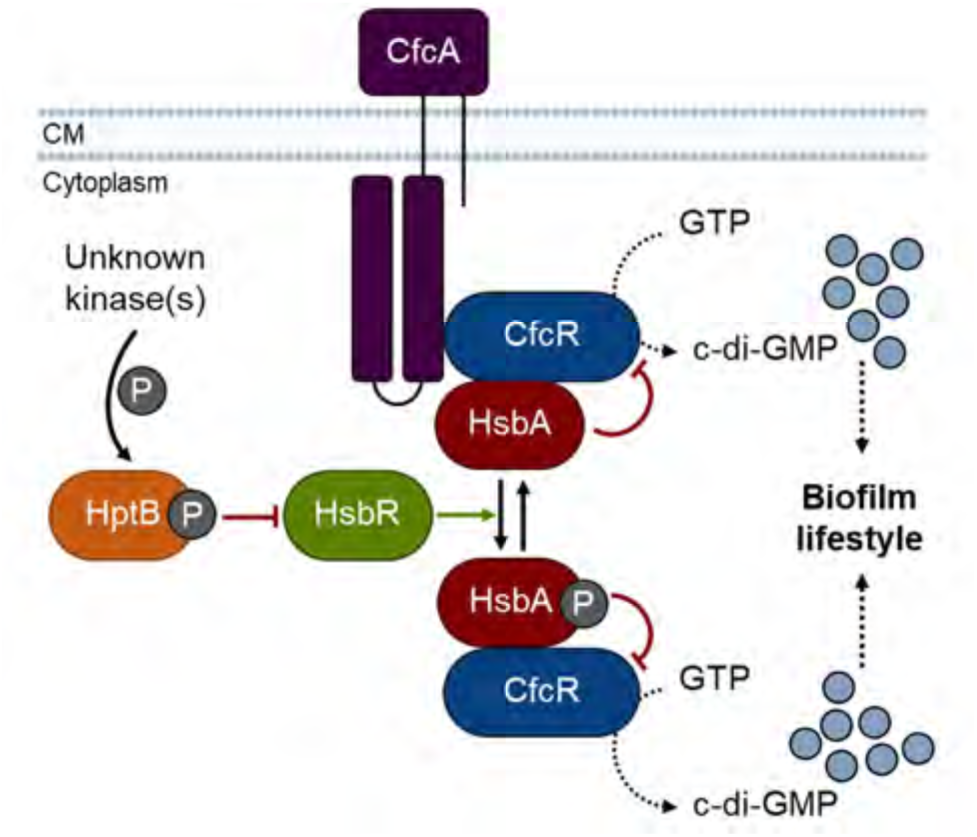
Working model of the regulation of biofilm formation by the HptB/Hsb/Cfc system. Green arrows and T-shaped end red lines denote positive and negative regulation. P: phosphoryl group. CM: cell membrane. Created in part with Biorender.com.

Our results also show that HsbA can inhibit biofilm formation in *P. putida* regardless of its phosphorylation state. This is evidenced by the fact that, unlike the Δ*hsbA* mutant, mutants lacking HptB or HsbR display wild-type biofilm phenotypes **(Fig. 5)**. In addition, both the non-phosphorylatable and phosphomimetic derivatives HsbA^S56A^ and HsbA^S56D^ proteins complement the biofilm-related phenotypes of the Δ*hsbA* mutant **(Fig. 6)**. However, under equivalent conditions HsbA^S56A^ fully restored expression of the c-di-GMP reporter pCdrA::*gfp*(ASV)^C^, while complementation by HsbA^S56D^ was partial, suggesting that phosphorylation may decrease the ability of HsbA to stimulate c-di-GMP synthesis. However, since this phenotype is not reproduced by the Δ*hptB* mutant, in which HsbA is expected to be phosphorylated, we favor the view that phosphorylated HsbA is no less efficient in stimulating the CfcR DGC activity than its unphosphorylated counterpart. Instead, we propose that the observed defect is a specific feature of the phosphomimetic mutant protein that does not fully replicate the activity of the native phosphorylated protein, a common observation in this type of mutants **(Klose *et al*., 1993, Horstmann *et al*., 2017)** Both *P. aeruginosa* HsbA^S56A^ and HsbA^S56D^ were able to stimulate biofilm formation and inhibit swarming motility, but only the effect of the HsbD-interacting HsbA^S56D^ was dependent on HsbD, suggesting that unphosphorylated HsbA may achieve the same regulatory outcome by a different, HsbD-independent mechanism **(Valentini *et al*., 2016)**. Since both forms of *P. putida* HsbA co-immunoprecipitate with CfcR **(Fig. S11)**, we propose that formation of an HsbA-CfcR complex inhibits CfcR DGC activity or CfcA-dependent phosphorylation of CfcR, regardless of the phosphorylation state of HsbA or the intracellular location of the complex **(Fig. 8)**. To provide further support to this proposal, we used AlphaFold to predict the possible interaction between CfcR and non-phosphorylated or phosphorylated HsbA **(Fig. S15)**. In both cases the output showed an interaction between monomeric HsbA and the intervening region between the GGDEF and EAL domains of CfcR, immediately upstream from the EAL domain **(Fig. S15C)**. Key interactions at the CfcR-HsbA contact interface occurred between the α-helix connecting the CfcR GGDEF and EAL domains, the first α-helix of the EAL domain, and the three α-helices of HsbA **(Fig. S15D)**. The location of the putative HsbA binding region supports a direct effect on the DGC activity, rather than on CfcA-depedent phosphorylation. Since the predicted binding site is in the vicinity of the EAL domain, we also considered the possibility that HsbA may induce a c-di-GMP PDE activity residing in this domain. However, sequence analyses performed by **Matilla *et al*. (2011)** revealed that critical conserved residues required for an active EAL domain **(Römling, 2009)** are missing, suggesting that CfcR cannot hydrolyze c-di-GMP. Because inactive EAL domains often play a regulatory role by modulating other activities of the protein or its interactions with other proteins or ligands **(Hengge *et al*., 2023)**, it is feasible that HsbA may act by disrupting some intra- or interdomain interaction key to such regulatory function. Although we acknowledge the limitations of computer-based modelling, the structural prediction is consistent with our observation that HsbA-CfcR interaction is independent of phosphorylation and in any case, it provides an excellent framework for further exploration aimed at clarifying the nature of the effect of HsbA on CfcR function.

Local nutrient limitation has also been shown to play an important role in biofilm maturation in different bacteria **(Runbaugh and Sauer, 2020)**. During biofilm growth, increasing microcolony size results in limited diffusion that leads to local nutrient depletion at the colony core. In these locations, groups of dispersing cells release themselves from the biofilm matrix, resume flagellar motility and eventually break through the colony outer cell layers, leaving behind hollow structures that eventually grow into the mature biofilm. This process, known as seeding dispersal **(Purevdorj-Gage *et al*., 2005)**, contributes to the three-dimensional remodeling of the biofilm structure that leads to mature biofilms **(Barken *et al*., 2008; Ma *et al*., 2009)**. Dispersing biofilm cells display unique phenotypes that enable them to resume the planktonic lifestyle, while being primed for recolonization at new sites **(Rollet *et al*., 2009; Chua *et al*., 2014; Runbaugh and Sauer, 2020)**. Seeding dispersal is well documented in *P. putida* **(Tolker-Nielsen *et al*., 2000; Langmead *et al*., 2006)**, and several lines of evidence suggest that dispersing cells likely play a role in shaping the mature biofilm architecture. Firstly, while early biofilm microcolonies are mostly clonal, mature biofilm macrocolonies display mixed populations, suggesting that dispersing cells may be reintegrated in the structure at random sites **(Tolker-Nielsen *et al*., 2000; Klausen *et al*., 2006)**. In addition, synthesis of some components of the biofilm matrix, notably the high molecular weight adhesin LapF and the extracellular polysaccharide Pea, is specifically induced by both the GSR and the SR **(Martínez-Gil *et al*., 2010; Liu *et al*., 2017)**. Our RNA-seq results confirm (p)ppGpp-dependent induction of *lapF*, its specific secretion system *lapHIJ*, and the *pea* polysaccharide cluster **(Fig. S1)**. We also detected enhanced transcription of the *csu* cluster, encoding orthologs of the components of the non-archetipal CupE chaperon-usher fimbriae, involved in micro- and macrocolony formation and maintenance of the three-dimensional architecture of the biofilm in *P. aeruginosa* **(Giraud *et al*., 2011; De Bemtzmann *et al*., 2012)**. Taken together, these observations suggest that local induction of the GSR and the SR results in local dissolution followed by reattachment of the dispersing cells, accompanied by qualitative changes in the composition of the mature biofilm matrix. On the other hand, while the SR is strictly activated by nutritional stress, the GSR is responsive to multiple forms of stress. It is therefore feasible that HsbAR-HptB and CfcR be synthesized in response to cues other than nutrient limitation **(Bouillet *et al*., 2024)**. Osmotic stress is one of the signals known to induce the GSR. Previous work showed that CfcR DGC is activated under high salt concentrations **(Tagua *et al*., 2022)**. It is tempting to speculate that salt stress may antagonize HsbA-dependent inhibition by an as of yet unknown mechanism to enable CfcR DGC activity.

The production of the DGC CfcR along with its cognate activator CfcA in response to nutrient limitation suggests the potential of inducing c-di-GMP synthesis at this stage. However, such potential is negated by the simultaneous production of the antagonist protein HsbA. We are currently exploring the hypothesis that regulation of CfcR DGC activity by HsbA is an integral part of the biofilm maturation process. In this scenario, local nutrient depletion within the microcolonies results in induction of the SR and the GSR and the subsequent synthesis of HsbA, HsbR, HptB and CfcR. While seeding dispersal occurs, HsbA efficiently prevents c-di-GMP synthesis by CfcR. However, as cells are released from the biofilm they encounter nutrient-sufficient conditions or other environmental cues that inactivate HsbA and release CfcR repression. This in turn provides an opportunity to terminate the dispersal program and reattach to resume biofilm development. We propose that the design of the system enables dispersing cells to be primed for rapid reinitiation of biofilm formation during the dispersal stage, provided that specific conditions (e.g., nutrient-sufficient conditions outside of the biofilm structure) are met.

The result of HsbA-dependent regulation of biofilm growth in *P. putida* is in sharp contrast with the observations previously reported in *P. aeruginosa*. In this organism, HsbA positively regulates biofilm formation by stimulating the activity of the DGC HsbD **(Valentini *et al*., 2016; Bouillet *et al*., 2019)**. So far, this regulation has not been linked to nutritional stress or stationary phase conditions, the SR or the GSR. We have examined the occurrence of *hsbA*, *hsbD* and *cfcR* orthologs in >1500 genomes in the *Pseudomonas* genome database (pseudomonas.com). Over 98% of the genomes analyzed contained an *hsbA* ortholog, suggesting that this signal transduction pathway is widespread in the genus. Over 97% of the HsbA^+^ strains carried at least an ortholog of *cfcR* or *hsbD*, and the co-occurrence rate of *hsbA* with *cfcR* or *hsbD* was 47% and 52%, with only 3% bearing both *cfcR* and *hsbD* **(Supplementary Data SD2)**. Orthologs of *hsbD* are largely restricted to the *P. aeruginosa* lineage, as previously described **(Valentini *et al*., 2016)**. In contrast, *cfcR* orthologs are widespread in the *P. fluorescens* lineage and the *P. stutzeri* group, and are only found sparingly in other members of the *P. aeruginosa* lineage. These results suggest that CfcR and HsbD are alternative targets to HsbA-dependent regulation that have been co-opted during the evolution of the genus *Pseudomonas* to achieve opposite outcomes in the regulation of biofilm growth. Bacteria thriving in soils are often subjected to one or more forms of environmental stress. *P. putida* has likely evolved to adapt to these conditions by promoting biofilm formation to colonize environments in which beneficial conditions prevail, such as the rhizosphere, and triggering biofilm dispersal in response to stress conditions, such as nutrient limitation **(Gjermanssen *et al.,* 2005)**, to enable dissemination and colonization of other niches. On the other hand, since biofilm growth offers protection against different forms of aggression, many bacteria form biofilms as a protective response when exposed to hostile conditions **(Davey and O’Toole, 2000; Jefferson *et al*., 2004; McDougald *et al*., 2011)**. *P. aeruginosa* is a pathogen that uses biofilm formation as a means to evade the host defenses and promote the development of a chronic infection, and therefore biofilm growth prevails under stress conditions **(Byrd *et al*., 2011; Tuon *et al*., 2022)**. *P. aeruginosa* biofilms disperse upon an upshift in carbon and energy source availability **(Sauer *et al*., 2004)**, although dispersal in response to other signals has also been documented **(Runbaugh and Sauer, 2020)**. Interestingly, the occurrence of separate mechanisms for biofilm formation in nutrient-sufficient and nutrient-limited conditions in *Shewanella putrefaciens* was recently documented **(Sun *et al*., 2025)**. Altogether, the work presented here provides a paramount example of how bacterial evolution can operate by rewiring different combinations of conserved and variable regulatory modules (phosphorelay cascades, two-component systems, effector proteins) to achieve different or even opposite regulatory outcomes to suit best their physiological requirements.

## ACKNOWLEDGMENTS

We wish to thank Rubén de Dios and Tamara Martín for their help with some of the constructions, Eloísa Andújar, from the Genomics Facility at CABIMER (Sevilla, Spain), Laura Tomás and Kathy García, from the Proteomics and Biochemistry and the Microscopy Facilities at CABD (Sevilla, Spain), and Ana Rodríguez-Hortal from the Mass Spectroscopy Facility at Universidad Pablo de Olavide for excellent technical assistance. This work was supported by grants PGC2018-097151-B-I00, PID2021-126121-NB-I00 and CEX2020-00108-M, funded by the Spanish Ministerio de Ciencia e Innovación/Agencia Española de Investigación (MCIN/AEI) and the European Regional Development Fund “A way of making Europe” (ERDF), and by a predoctoral Formación de Profesorado Universitario contract 19/02899 of the Spanish Ministerio de Educación y Formación Profesional, awarded to M P-S. Funding for open access publishing provided by Universidad Pablo de Olavide/CBUA.

