## Supplemental Materials for "HsbA represses stationary phase biofilm formation in *Pseudomonas putida*"

### SUPPLEMENTARY INFORMATION

#### 1. SUPPLEMENTARY MATERIALS AND METHODS

##### Plasmid and strain construction

**Construction of MRB171, MRB178, MRB179, MRB193, MRB229, MRB230, MRB231 and MRB232.** To construct *P. putida* in-frame deletion mutants of the *hsbA, hptB, hsbR, rpoS, cfcR* and *cfcA*, upstream and downstream chromosomal regions flanking these genes were PCR-amplified using appropriate oligonucleotide pairs specified in **Table S1**. Upstream and downstream *hsbA, cfcR* and *cfcA* PCR products were cleaved with EcoRI and BamHI (upstream region) or BamHI and XbaI (downstream region), and three-way ligated into EcoRI- and XbaI-digested gene replacement vector pEMG that incorporates two I-SceI sites flanking the *lacZ* polylinker, yielding pMR297, pMRB445 and pMRB446, respectively. Upstream and downstream *hptB* and *hsbR* PCR products were cleaved with SacI and BamHI (upstream region) or BamHI and XbaI (downstream region), and three-way ligated into SacI- and XbaI-digested pEMG, yielding pMRB318 and pMRB320, respectively. Upstream and downstream *rpoS* PCR products were assembled by overlap extension PCR and cloned into SmaI-digested pEMG, yielding pMR359. These plasmids were transferred by triparental mating to *P. putida* KT2442 harboring the pSW-I plasmid, expressing I-SceI from the 3-methyl benzoic acid-inducible *xylS*-P*m* system. Selection of integration and allelic replacement by homologous recombination-based repair of the chromosomal cleavage at the integrated I-SceI site was performed as described **(Martínez-García & de Lorenzo, 2011)** to generate Δ*hsbA* (MRB171)*,* Δ*hptB* (MRB178)*,* Δ*hsbR* (MRB179)*,* Δ*rpoS* (MRB193)*,* Δ*cfcR* (MRB229) and Δ*cfcA* (MRB230) mutant strains, respectively. Similarly, pMRB445 and pMRB446 were transferred to Δ*hsbA* to generate Δ*hsbA*Δ*cfcR* (MRB231) and Δ*hsbA*Δ*cfcA* (MRB232) double mutants. Loci deletions were verified by PCR. Curation of pSW-I plasmid was carried out by successive growth cycles in LB without antibiotics.

**Construction of pMRB397, pMRB475 and pMRB474.** For HsbA-GFP and CfcA-GFP expression under their natural P*hsbA* or P*cfcA* promoters, P*hsbA-hsbA* or P*cfcA-cfcA* chromosomal regions were PCR-amplified and cloned into SpeI- and BamHI-digested pMRB187, yielding pMRB397 and pMRB475, respectively. For CfcR-GFP expression under its natural P*cfcR* promoter, a P*cfcR-cfcR* chromosomal region was PCR-amplified and cloned into SpeI- and XmaI-digested pMRB187, yielding pMRB474.

**Construction of pMRB369, pMRB388 and pMRB418.** For HsbA expression under the P*sal* promoter, a pMRB172-derivative was constructed by cloning the PCR-amplified *hsbA* ORF into the SpeI- and PstI-digested pMRB172, yielding pMRB369. For $HsbA^{S56A}$ and $HsbA^{S56D}$ expression under the P*sal* promoter, site-directed mutagenesis PCR was performed for serine-to-alanine or serine-to-aspartic acid replacement with appropriate oligonucleotides using pMRB369 as DNA template, and overlap extension PCR products holding P*sal*-$hsbA^{S56A}$ or P*sal*-$hsbA^{S56D}$ were cloned into SpeI- and PstI-digested pMRB172, yielding pMRB388 and pMRB418, respectively.

**Construction of pMRB417, pMRB416, pMRB415, pMRB438 and pMRB439.** For HsbA expression under its natural P*hsbA* promoter, the *hsbA* ORF from pMRB397 was PCR-amplified, and a SalI-PstI 175 bp fragment was cloned into pMRB397 digested with the same enzymes, yielding pMRB417. For $HsbA^{S56A}$ and $HsbA^{S56D}$ expression under their natural P*hsbA* promoter, pMRB388 and pMRB418 were SalI- and PstI-digested, and the 175 bp fragments were cloned into pMRB397 digested with the same enzymes, yielding pMRB416 and pMRB415, respectively. For $HsbA^{S56A}$-GFP and $HsbA^{S56D}$-GFP expression under their natural P*hsbA* promoter, a 1631 bp fragment derived from EcoRV-digested pMRB416 or pMRB415, was cloned into the 4133 bp fragment derived from EcoRV-digested pMRB397, yielding pMRB438 and pMRB439, respectively.

**Co-immunoprecipitation assays**

To identify interactors of HsbA, $HsbA^{S56A}$ and $HsbA^{S56D}$, we used strains bearing miniTn7 transposons to produce GFP-tagged bait proteins from the P*hsbA* promoter, or solely GFP from the constitutive P*A1/04/03* promoter as negative control. The cells were grown to an $OD_{600}$ of 4.0 in 100 mL LB medium and harvested by centrifugation (5000 *g*, 4 ºC, 15 min). Cells were washed three times with cold 10 mM HEPES buffer, pelleted and stored at -80 ºC for further processing. Pelleted cells were resuspended in 10 mL cold solubilization buffer (20 mM Tris-HCl pH 8.0, 150 mM NaCl, 0.5% Nonidet P-40) containing a protease inhibitor cocktail (1 mM phenylmethylsulfonyl fluoride [PMSF], 10 mg $l^{-1}$ leupeptin, 10 mg $l^{-1}$ pepstatin) and were disrupted by sonication. Lysates were clarified by centrifugation (13000 *g*, 4 ºC, 5 min) and soluble fractions were rotating-incubated overnight at 4 ºC with 20 µL of GFP-Trap® agarose beads (ChromoTek) equilibrated with solubilization buffer. Beads were then collected by centrifugation (2500 *g*, 4 ºC, 5 min) and washed ten times in 1 mL solubilization buffer. Proteins were eluted by resuspending the beads in 100 µL 2x SDS-loading buffer (150 mM Tris-HCl pH 6.8, 20% glycerol, 4% SDS, 0.1% bromophenol blue, 10% β-mercaptoetanol) and boiled at 100 ºC for 10 min. Soluble fractions were collected after centrifugation (13000 *g*, 4 ºC, 1 min), and proteins were precipitated by addition of 25 µL cold trichloroacetic acid followed by 10 min of ice-incubation. Pellets were washed three times with 200 µL cold acetone and dried at 95 ºC for 10 min for storage at 4 ºC. Protein identification by nLC-MS/MS was carried out at the BIO-MS facility from Pablo de Olavide University (Biomolecular Mass Spectrometry, Proteomics and Metabolomics Laboratory, Sevilla, Spain). Protein pellets were resuspended in 100 µL buffer A (6 M urea, 50 mM ammonium bicarbonate, 10 mM dithiothreitol [DTT]) and incubated for 60 min at room temperature. Samples were next added 30 mM iodoacetamide (IAA) and incubated for 30 min in the dark. Samples were digested at 37 ºC overnight using bovine trypsin (Sequencing Grade Modified Trypsin, Promega) in a trypsin-to-protein ratio 1:12 (w/w) and reactions were stopped by addition of 0.5% formic acid. Bond Elute OMIX C18 tips (Agilent

Technologies) were used to concentrate and desalt peptide extracts, following the manufacturer's instructions. Samples were dried under vaccum and eluted in 0.1% trifluoroacetic acid. Peptide samples (5 µg) were separated in an EASY-nLC™ system (Thermo Scientific) using a 50 cm C18 EASY-Spray™ HPLC column (Thermo Scientific). The following solvents were used as mobile phases: water with 0.1% formic acid (phase A) and 80% acetonitrile, 20% water with 0.1% formic acid (phase B). Separation was achieved with an acetonitrile gradient: 10-35% phase B over 120 min, 35-100% phase B over 1 min, and 100% phase B over 5 min at a flow rate of 200 nL/min. A Q Exactive™ Plus Orbitrap™ mass spectrometer (Thermo Scientific) was used to acquire the top 10 MS/MS spectra in data-dependent acquisition (DDA) mode. LC-MS data were analyzed using the SEQUEST® HT search engine from Thermo Scientific™ Proteome Discoverer™ 2.2 software. Data were searched against the UniProt *Pseudomonas putida* KT2440 protein database and results were filtered using a 1% protein FDR threshold. Peptide count data was extracted and one unit was added to all counts to remove the zeroes. Differential protein abundance was calculated as the peptide counts ratio between the sample and the negative control. Two separate replicates of each strain were analyzed in this assay.

#### 2. SUPPLEMENTARY TABLE

**Table S1. Bacterial strains, plasmids and oligonucleotides used in this work.** Cm: chloramphenicol. Rif: rifampicin. Km: kanamycin. Ap: ampicillin. Gm: gentamycin. Tel: tellurite.

| Bacterial strain | Genotype/phenotype | Reference/source |
|---|---|---|
| ***E. coli*** | | |
| DH5α | Φ80d*lacZ*ΔM15 Δ(*lacZYA-argF*)U169 *recA*1 *endA*1 *hsdR*17 ($r_k^-$ $m_k^+$) *supE*44 *thi*-1 *gyrA* *relA*1 | Hanahan, 1983 |
| DH5α λpir | DH5α with lysogenic phage λ-pir, host for R6K replication origin plasmids | Víctor de Lorenzo |
| ***P. putida*** | | |
| KT2440 | mt-2 *hsdR*1 ($r^-$ $m^+$) | Franklin *et al*., 1981 |
| KT2440tel | KT2440 tagged with miniTn5-Tel. Tel$^r$ | Sze *et al*., 2002 |
| KT2442 | mt-2 *hsdR*1 ($r^-$ $m^+$). Cm$^r$ Rif$^r$. | Franklin *et al*., 1981 |
| MRB171 | KT2442 Δ*hsbA*. Cm$^r$ Rif$^r$ | This work |
| MRB172 | KT2442 Δ*flgM*. Cm$^r$ Rif$^r$. | Pulido-Sánchez *et al.,* 2025 |
| MRB176 | KT2442 Δ*fliA*. Cm$^r$ Rif$^r$. | Pulido-Sánchez *et al.,* 2025 |
| MRB178 | KT2442 Δ*hptB*. Cm$^r$ Rif$^r$ | This work |
| MRB179 | KT2442 Δ*hsbR*. Cm$^r$ Rif$^r$ | This work |
| MRB193 | KT2442 Δ*rpoS*. Cm$^r$ Rif$^r$ | This work |
| MRB229 | KT2442 Δ*cfcR*. Cm$^r$ Rif$^r$ | This work |
| MRB230 | KT2442 Δ*cfcA*. Cm$^r$ Rif$^r$ | This work |
| MRB231 | KT2442 Δ*hsbA*Δ*cfcR*. Cm$^r$ Rif$^r$ | This work |
| MRB232 | KT2442 Δ*hsbA*Δ*cfcA*. Cm$^r$ Rif$^r$ | This work |
| MRB52 | KT2442 Δ*fleQ*. Cm$^r$ Rif$^r$. | Navarrete *et al*., 2019 |
| MRB62 | KT2442 *fliP*::miniTn5-Km [fT3SS$^-$]. Cm$^r$ Rif$^r$ Km$^r$ | López-Sánchez *et al*., 2016 |
| PP1922 | KT2440tel Δ*relA*::Km Δ*spoT*::Gm [ppGpp$^0$]. Tel$^r$ Km$^r$ Gm$^r$ | Díaz-Salazar *et al.,* 2017 |

| Plasmid | Genotype/phenotype | Reference/source |
|---|---|---|
| pBK-miniTn7-ΩGm | pUC19-based delivery plasmid for miniTn7-ΩGm. Mob$^+$, Ap$^r$ Gm$^r$ | Koch *et al.,* 2001 |
| pBK-miniTn7-*gfp*2 | pBK-miniTn7-ΩGm-derived vector containing a P*A1/04/03*-*gfp*mut3 transcriptional fusion. Mob$^+$, Ap$^r$ Gm$^r$ | Lambertsen *et al.,* 2004 |
| pCdrA::*gfp*(ASV)$^C$ | pUCP22Not-P*cdrA*-RBSII-*gfp*ASV-T0-T1. Ap$^r$ Gm$^r$ | Rybtke *et al*., 2012 |
| pEMG | pJP5603 bearing a *lacZ*α polylinker with two flanking I-SceI sites. R6K, Mob$^+$, Km$^r$ | Martínez-García & de Lorenzo, 2011 |
| pMRB1 | pBBR1-MCS4-derived broad host-range *gfp*mut3::*lacZ* transcriptional fusion vector. pBBR1, Mob$^+$, Ap$^r$ | Jiménez-Fernández *et al*., 2015 |
| pMRB172 | pUC18Sfi-miniTn*7*BB-Gm-based delivery plasmid for miniTn*7*BB-Gm [*nahR*-P*sal*]. Ap$^r$ Gm$^r$ | Leal-Morales *et al*., 2022 |
| pMRB187 | pUC18Sfi-miniTn*7*BB-Gm-based delivery plasmid for C-terminal *gfp*-mut3 translational fusions. Ap$^r$ Gm$^r$ | Pulido-Sánchez *et al.,* 2025 |
| pMRB267 | pMRB3-derived vector containing a *gfp*mut3::*lacZ* transcriptional fusion to the P*hsbA* promoter. Ap$^r$ | Jiménez-Fernández *et al*., 2016 |
| pMRB276 | pMRB3-derived vector containing a *gfp*mut3::*lacZ* transcriptional fusion to the P*cheA* promoter. Ap$^r$ | Jiménez-Fernández *et al*., 2016 |
| pMRB297 | pEMG bearing a 1189 bp insert with upstream and downstream flanking regions of *hsbA*. Km$^r$ | This work |
| pMRB3 | pMRB1-derived vector containing the Gateway conversion cassette *attR2*-*ccdB*-Cm$^r$-*attR1*. Ap$^r$ Cm$^r$ | Jiménez-Fernández *et al*., 2014 |
| pMRB318 | pEMG bearing a 1113 bp insert with upstream and downstream flanking regions of *hptB*. Km$^r$ | This work |
| pMRB320 | pEMG bearing a 1097 bp insert with upstream and downstream flanking regions of *hsbR*. Km$^r$ | This work |
| pMRB359 | pEMG bearing a 2123 bp insert with upstream and downstream flanking regions of *rpoS*. Km$^r$ | This work |
| pMRB369 | pMRB172-derived vector containing *nahR*-P*sal*-*hsbA*. Ap$^r$ Gm$^r$ | This work |
| pMRB388 | pMRB172-derived vector containing *nahR*-P*sal*-*hsbA*$^{S56A}$. Ap$^r$ Gm$^r$ | This work |
| pMRB397 | pMRB187-derived vector containing P*hsbA*-*hsbA* translationally fused to *gfp*mut3 at its C-terminus. Ap$^r$ Gm$^r$ | This work |

| | | |
|---|---|---|
| pMRB415 | pUC18Sfi-miniTn*7*BB-Gm-derived vector containing P*hsbA*-*hsbA*$^{S56D}$. Ap$^r$ Gm$^r$ | This work |
| pMRB416 | pUC18Sfi-miniTn*7*BB-Gm-derived vector containing P*hsbA*-*hsbA*$^{S56A}$. Ap$^r$ Gm$^r$ | This work |
| pMRB417 | pUC18Sfi-miniTn*7*BB-Gm-derived vector containing P*hsbA*-*hsbA*. Ap$^r$ Gm$^r$ | This work |
| pMRB418 | pMRB172-derived vector containing *nahR*-P*sal*-*hsbA*$^{S56D}$. Ap$^r$ Gm$^r$ | This work |
| pMRB438 | pMRB187-derived vector containing P*hsbA*-*hsbA*$^{S56A}$ translationally fused to *gfp*mut3 at its C-terminus. Ap$^r$ Gm$^r$ | This work |
| pMRB439 | pMRB187-derived vector containing P*hsbA*-*hsbA*$^{S56D}$ translationally fused to *gfp*mut3 at its C-terminus. Ap$^r$ Gm$^r$ | This work |
| pMRB445 | pEMG bearing a 1081 bp insert with upstream and downstream flanking regions of *cfcR*. Km$^r$ | This work |
| pMRB446 | pEMG bearing a 1047 bp insert with upstream and downstream flanking regions of *cfcA*. Km$^r$ | This work |
| pMRB474 | pMRB187-derived vector containing P*cfcR*-*cfcR* translationally fused to *gfp*mut3 at its C-terminus. Ap$^r$ Gm$^r$ | This work |
| pMRB475 | pMRB187-derived vector containing P*cfcA*-*cfcA* translationally fused to *gfp*mut3 at its C-terminus. Ap$^r$ Gm$^r$ | This work |
| pRK2013 | Helper plasmid for triparental mating. ColE1, Mob$^+$, Km$^r$ | Figurski & Helinski, 1979 |
| pSW-I | Plasmid expressing I-SceI from *xylS*-P*m*. RK2, Mob$^+$, Ap$^r$ | Wong & Mekalanos, 2000 |
| pTNS2 | Helper plasmid expressing the *Tn*7 transposase. R6K, Mob$^+$, Ap$^r$ | Choi *et al*., 2005 |
| pUC18Sfi-miniTn*7*BB-Gm | pUC18SfiI-based delivery plasmid for the synthetic minitransposon mini*Tn*7BB-Gm. Ap$^r$ Gm$^r$ | Jiménez-Fernández *et al*., 2014 |

| **Oligonucleotide** | **Sequence (5' to 3')** | **Use** |
|---|---|---|
| CfcA-BamHI_rev | CAGTGGATCCAATTCGCTCCAGTTGCGG | P*cfcA* promoter and *cfcA* ORF amplification |
| CfcA-DOWN-BamHI_fwd | CAGTGGATCCCTGTTCTCGTTGATCCGTG | *cfcA* downstream region amplification |
| CfcA-DOWN-DOWN_rev | AGTACTCGGCGAAGTCACG | *cfcA* chromosomal deletion verification |
| CfcA-DOWN-XbaI_rev | ACTGTCTAGAGTAGATGCCCTGCTTGGC | *cfcA* downstream region amplification |
| CfcA-qRT-PCR_fwd | GCAGCATGGTCGAGGACAAC | qRT-PCR of *cfcA* ORF |
| CfcA-qRT-PCR_rev | CTCGTTACCGAAGTCGTTCCA | qRT-PCR of *cfcA* ORF |
| CfcA-UP-BamHI_rev | ACTGGGATCCCATGAGAAAACGCCCGAG | *cfcA* upstream region amplification |
| CfcA-UP-EcoRI_fwd | CAGTGAATTCGATCAAGTGCATGCCCTC | *cfcA* upstream region amplification |
| CfcA-UP-UP_fwd | GAATACCCACCACCGTGTC | *cfcA* chromosomal deletion verification |
| CfcR-DOWN-DOWN_rev | ATCGATCAGCCAGGCCAC | *cfcR* chromosomal deletion verification |
| CfcR-DOWN-BamHI_fwd | CAGTGGATCCTGATTGGCTAGAATCGGC | *cfcR* downstream region amplification |
| CfcR-DOWN-XbaI_rev | ACTGTCTAGAACTTCCATGCTGCATGTTC | *cfcR* downstream region amplification |
| CfcR-qRT-PCR_fwd | CTTACCCTCAACAGCAGCCAC | qRT-PCR of *cfcR* ORF |
| CfcR-qRT-PCR_rev | GAGCTTGCCGTCAATACTTGC | qRT-PCR of *cfcR* ORF |
| CfcR-UP-BamHI_rev | CATGGGATCCCATGCTGCTTCTCTGTTATGTG | *cfcR* upstream region amplification |
| CfcR-UP-EcoRI_fwd | CATGGAATTCGCGATCATCAACCAGTACG | *cfcR* upstream region amplification |
| CfcR-UP-UP_fwd | GATGCACGGTTCGTCCTC | *cfcR* chromosomal deletion verification |
| CfcR-XmaI_rev | CAGTCCCGGGCGGCCCTGGCCAGGAGAA | P*cfcR* promoter and *cfcR* ORF amplification |
| HptB-DOWN-BamHI_fwd | GATCGGATCCTGATACGCGAGTTGGCG | *hptB* downstream region amplification |
| HptB-DOWN-DOWN_rev | GACGGCTCTTCAGGCAGT | *hptB* chromosomal deletion verification |
| HptB-DOWN-XbaI_rev | GATCTCTAGAAAACCTGGGTTTGCGCC | *hptB* downstream region amplification |
| HptB-qRT-PCR_fwd | CAGCTCACAGTTTCAAGG | qRT-PCR of *hptB* ORF |
| HptB-qRT-PCR_rev | ATCAAGTCTTCAATACCGTAC | qRT-PCR of *hptB* ORF |
| HptB-UP-BamHI_rev | GATCGGATCCCACTTGTTCACTCCTTGATCA | *hptB* upstream region amplification |
| HptB-UP-SacI_fwd | GATCGAGCTCCTGAGCTTCGAGGTGCGT | *hptB* upstream region amplification |
| HptB-UP-UP_fwd | TCTGATAGTGGCCGCTCC | *hptB* chromosomal deletion verification |
| HsbA-BamHI_rev | CATGGGATCCACTGATATCGAAGAGTTTTTCGAAATTG | P*hsbA* promoter and *hsbA* ORF amplification |
| HsbA-deP_fwd | GACACCACCTACCTCGACGCCTCCGCGCTCGGCATGCTG | *hsbA* ORF amplification with S56A replacement |
| HsbA-deP_rev | CAGCATGCCGAGCGCGGAGGCGTCGAGGTAGGTGGTGTC | *hsbA* ORF amplification with S56A replacement |

| | | |
|---|---|---|
| HsbA-DOWN-BamHI_fwd | CTCCGGATCCCGCAAGATTCTCGCCATC | *hsbA* downstream region amplification |
| HsbA-DOWN-DOWN_rev | AATGCCGCCAGCATCAGG | *hsbA* chromosomal deletion verification |
| HsbA-DOWN-XbaI_rev | TGAATCTAGAGTAAGGCGATTGCAGGTAGC | *hsbA* downstream region amplification |
| HsbA-P_fwd | GACACCACCTACCTCGACGCCTCCGCGCTCGGCATGCTG | *hsbA* ORF amplification with S56D replacement |
| HsbA-P_rev | CAGCATGCCGAGCGCGGAGGCGTCGAGGTAGGTGGTGTC | *hsbA* ORF amplification with S56D replacement |
| HsbA-PstI_rev | GCATCTGCAGCGGCCGCTACTAGTATTATTAACTGATATCGAAGAGTTTTTCGA | *hsbA* ORF amplification |
| HsbA-qRT-PCR_fwd | CTTCGATTTCGGCAAGCATCAG | qRT-PCR of *hsbA* ORF |
| HsbA-qRT-PCR_rev | GAGGTAGGTGGTGTCCTTCAG | qRT-PCR of *hsbA* ORF |
| HsbA-UP-BamHI_rev | TCTCGGATCCCATGCTAGCTAGCGATTCCTTC | *hsbA* upstream region amplification |
| HsbA-UP-EcoRI_fwd | GGCGGAATTCGTGGACATGGCCGAAGAG | *hsbA* upstream region amplification |
| HsbA-UP-UP_fwd | CCGGCAGGTTGATAAGCC | *hsbA* chromosomal deletion verification |
| HsbA-XbaI_fwd | ATGCTCTAGAGAAAGAGGAGAAATACTAGATGGCAGTCGAGACTGATTTTTC | *hsbA* ORF amplification |
| HsbR-DOWN-BamHI_fwd | GATCGGATCCTGAGCCACGGGTTGGCT | *hsbR* downstream region amplification |
| HsbR-DOWN-DOWN_rev | CGAACGCGAGGTTTTGG | *hsbR* chromosomal deletion verification |
| HsbR-DOWN-XbaI_rev | GATCTCTAGACTTGCAGCAAAGGGTTGG | *hsbR* downstream region amplification |
| HsbR-qRT-PCR_fwd | AATGAAGAAGAGGGGCTG | qRT-PCR of *hsbR* ORF |
| HsbR-qRT-PCR_rev | GTCACGCTGTTCCAGTAC | qRT-PCR of *hsbR* ORF |
| HsbR-UP-BamHI_rev | GATCGGATCCCATGTCGGCGCTCAACTG | *hsbR* upstream region amplification |
| HsbR-UP-SacI_fwd | GATCGAGCTCCTCTATAGGTGTTCGCCGCAG | *hsbR* upstream region amplification |
| HsbR-UP-UP_fwd | TTGCTGGATGAGCTGTCG | *hsbR* chromosomal deletion verification |
| PcfcA-SpeI_fwd | CAGTACTAGTCGGCACATTGAAACTGTCG | P*cfcA* promoter and *cfcA* ORF amplification |
| PcfcR-SpeI_fwd | CAGTACTAGTGAAGGCATCAAGCACGGCG | P*cfcR* promoter and *cfcR* ORF amplification |
| PhsbA-SpeI_fwd | CATGACTAGTGGCACCAGAACAACCTCAATAAC | P*hsbA* promoter and *hsbA* ORF amplification |
| RpoS-DOWN-DOWN_rev | CCCGCTGTCACATGAGCAC | *rpoS* chromosomal deletion verification |
| RpoS-DOWN_fwd | CTCAGTAAAGAAGTGCCGGAGGAAAAGAATGGTCTGTCCAGC | *rpoS* downstream region amplification |
| RpoS-DOWN_rev | GGCTTGATGCGTACTTCTGC | *rpoS* downstream region amplification |
| RpoS-UP_fwd | GTCTTTTGTACCCGTCTGCC | *rpoS* upstream region amplification |
| RpoS-UP_rev | CTCCGGCACTTCTTTACTGAG | *rpoS* upstream region amplification |
| RpoS-UP-UP_fwd | GTGCCGCTGCTCAATGGTC | *rpoS* chromosomal deletion verification |
| Tn7-GlmS | AATCTGGCCAAGTCGGTGAC | Confirmation of miniTn7-derivatives integration |
| Tn7-R109 | CAGCATAACTGGACTGATTTCAG | Confirmation of miniTn7-derivatives integration |

#### 3. SUPPLEMENTARY DATA

**Supplementary Data SD1. Proteins identified in the co-immunoprecipitation analysis of HsbA, HsbA$^{S56A}$ and HsbA$^{S56D}$.** Tables provide the annotations, Uniprot accession numbers, peptide counts, enrichment scores ($log_2$ fold-change compared to the negative control) and p-values of all proteins co-inmmunoprecipitated with HsbA-GFP, HsbA$^{S56A}$-GFP and HsbA$^{S56D}$-GFP. Mass spectrometry raw data include the identified peptides and corresponding molecular weight.

**Supplementary Data SD2. Occurrence of HsbA, CfcR and HsbD orthologs in *Pseudomonas* genomes.** Orthologs were identified using Diamond PBLAST at the *Pseudomonas* genome database (pseudomonas.com) and the protein sequences of PP_4364 (HsbA), PP_4959 (CfcR) and PA3342 (HsbD) as seeds. The table displays the genomes bearing orthologs for all three proteins (green), HsbA and CfcR (blue), HsbA and HsbD (orange), or only HsbA (grey). Twenty-five genomes did not contain an HsbA ortholog, or the ortholog was a pseudogene, and were discarded from the analysis. The frequency of occurrence and co-occurrence of the three proteins is shown in columns P to R.

#### 4. SUPPLEMENTARY FIGURES

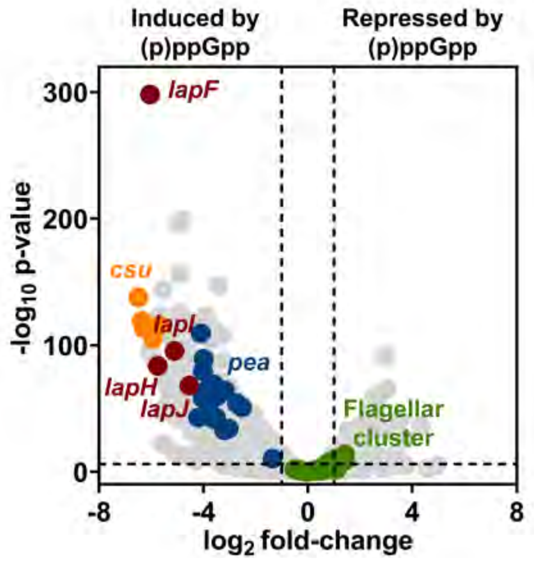


**Figure S1. Differential gene expression in the ppGpp$^0$ mutant compared to the wild-type strain.** Volcano plot of RNA-seq expression data (GEO:GSE281821) comparing differential gene expression of a ppGpp$^0$ mutant relative to the wild-type (KT2440) strain. The horizontal dotted line corresponds to an adjusted p-value of $10^{-6}$. Vertical dotted lines correspond to a twofold-change expression threshold. Coloured points highlight: the high molecular weight adhesin-encoding gene *lapF* and its associated ABC transporter *lapHIJ* (red), genes *pp2357-2363* encoding structural components of the type I Csu pili (orange), the *P. putida* exopolysaccharide a (Pea) biosynthesis gene cluster *pp3126-pp3142* (blue), and the flagellar gene cluster except *hsbA, hsbR* and *hptB* (green).

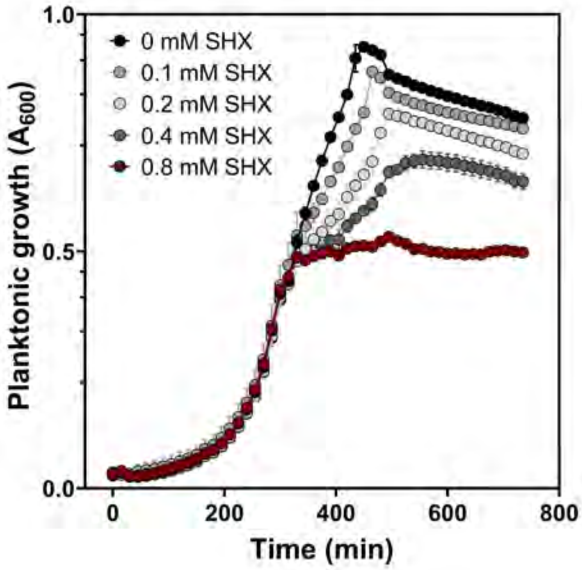


**Figure S2. Effect of serine hydroxamate on wild-type *P. putida* growth.** Planktonic growth curves of *P. putida* KT2440 in minimal medium. Serine hydroxamate (SHX) was added at an $OD_{600}$ of 0.4 at different concentrations. Points and error bars represent averages and standard deviations of at least three biological replicates.

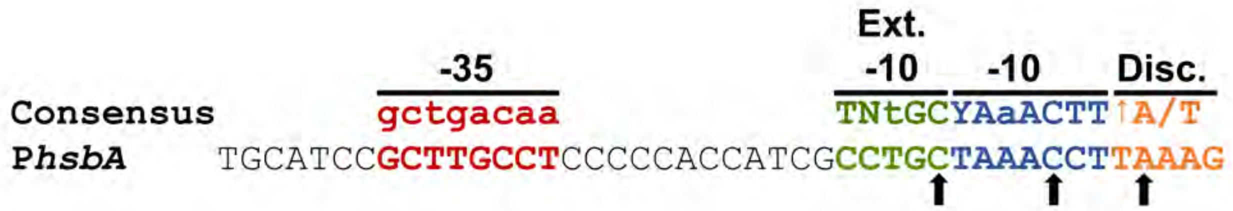


**Figure S3. Putative RpoS-dependent P*hsbA* promoter.** Sequence of the RpoS-consensus binding sequence in the region upstream from *hsbA*, showing the location of the -35, -10, extended -10 (Ext. -10) and discriminator (Disc.) Arrows indicate nucleotide positions of the discriminating -5A, -9C and -13C nucleotide.

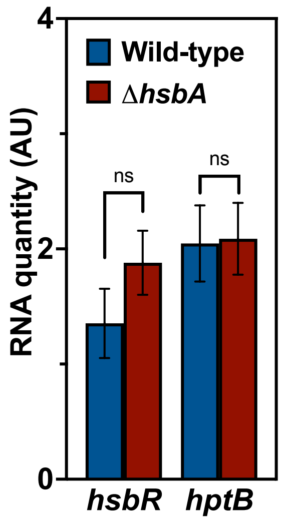


**Figure S4. Expression of *hsbR* and *hptB* in the Δ*hsbA* mutant.** qRT-PCR quantification of *hsbR* and *hptB* expression in wild-type (KT2442) and Δ*hsbA* cells harvested in stationary phase. Columns and error bars represent the averages and standard deviations of at least three biological replicates. Stars designate p-values for the Student's t-test for unpaired samples not assuming equal variance. ns: non-significant.

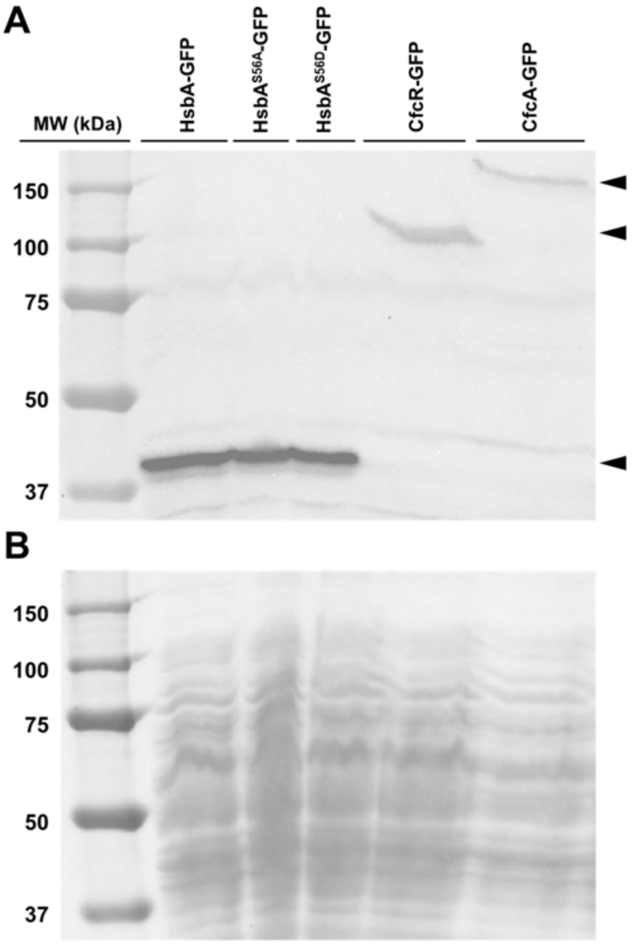


**Figure S5. Western blot of the HsbA-GFP, CfcR-GFP and CfcA-GFP fusion proteins.** Soluble extracts from the wild-type strain KT2442 bearing the P*hsbA-hsbA-gfp*, P*hsbA-hsbA*$^{S56A}$*-gfp*, P*hsbA-hsbA*$^{S56D}$*-gfp*, P*cfcR-cfcR-gfp* and P*cfcA-cfcA-gfp* transposons were resolved by SDS-PAGE and blotted to a nitrocellulose membrane. **A.** Western blot using anti-GFP antiserum. **B.** Ponceau S stain of the blotted membrane. The molecular weights of the visible size marker bands are indicated. Arrowheads denote bands corresponding to the HsbA-GFP (bottom), CfcR-GFP (middle) and CfcA-GFP (top) fusion proteins.

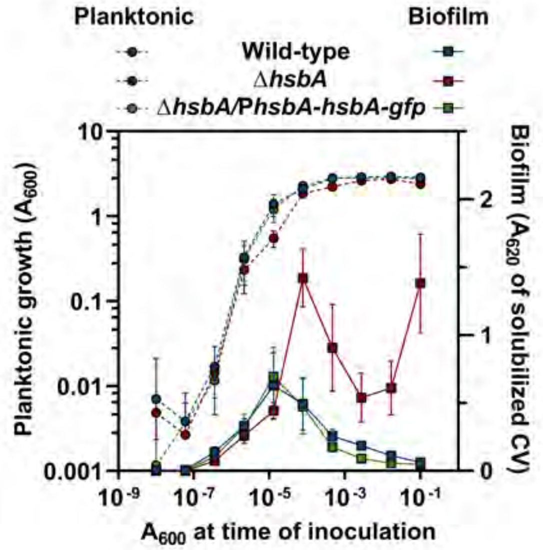


**Figure S6. Complementation of the Δ*hsbA* mutant.** Dilution series-based growth curves showing complementation of the Δ*hsbA* mutant with the P*hsbA-hsbA-gfp* transposon. Curves of the wild-type and Δ*hsbA* strains performed in parallel are shown as controls. Planktonic (left axis, circles) and biofilm growth (right axis, squares) are plotted against the initial $A_{600}$ of each dilution. CV: crystal violet. Points and error bars represent averages and standard deviations of six technical replicates. The plot shows a representative assay out of at least three biological replicates.

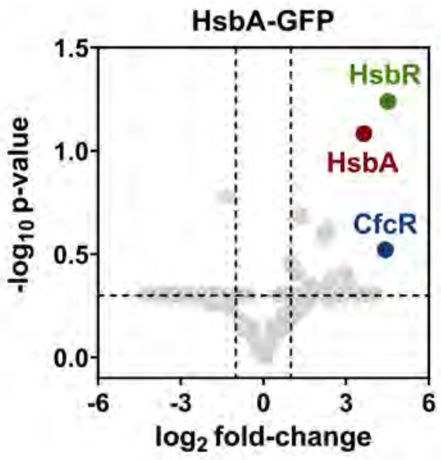


**Figure S7. Interactors of HsbA.** Volcano plot showing the proteins co-immunoprecipitated with HsbA-GFP in stationary-phase wild-type cells bearing the P*hsbA-hsbA-gfp* transposon, as identified by mass spectrometry. Points show the $log_2$ of the average fold-change in the peptide counts for each identified protein compared the negative control *vs.* the $-log_{10}$ p-value of these peptide counts (n=2). Coloured dots indicate proteins assessed in this work.

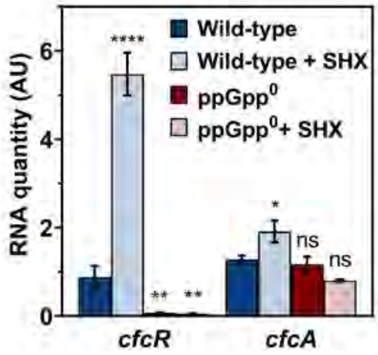


**Figure S8. Induction of the expression of *cfcR* and *cfcA* by serine hydroxamate.** qRT-PCR quantification of *cfcR* and *cfcA* expression in wild-type (KT2440) and ppGpp$^0$ cells harvested in exponential phase 2 h after the addition of 0.8 mM SHX in minimal medium. Columns and error bars represent the averages and standard deviations of at least three biological replicates. Stars designate p-values for the Student's t-test for unpaired samples not assuming equal variance. ns: non-significant; *:p<0.05; **:p<0.01; ***:p<0.001; ****:p<0.0001.

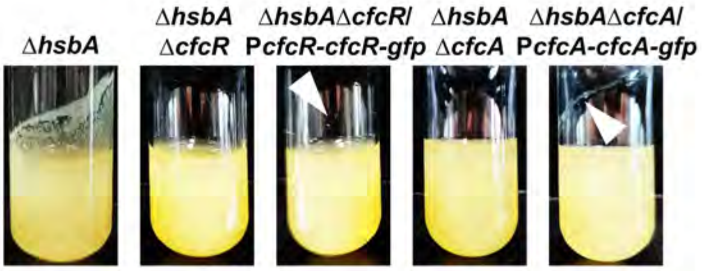

**Figure S9. Complementation of the Δ*hsbA*Δ*cfcR* and Δ*hsbA*Δ*cfcA* mutants.** Pellicle formation in glass tubes of LB cultures, showing complementation of the Δ*hsbA*Δ*cfcR* and Δ*hsbA*Δ*cfcA* mutants with the P*cfcR-cfcR-gfp* and P*cfcA-cfcA-gfp* transposons, respectively, after 48 h incubation. Cultures of the Δ*hsbA,* Δ*hsbA*Δ*cfcR* and Δ*hsbA*Δ*cfcA* strains are shown for comparison. Arrowheads denote formation of a thin pellicle in the complemented strain that is not present in the Δ*hsbA*Δ*cfcR* or Δ*hsbA*Δ*cfcA* mutants. Each picture shows a representative assay out of at least three biological replicates.

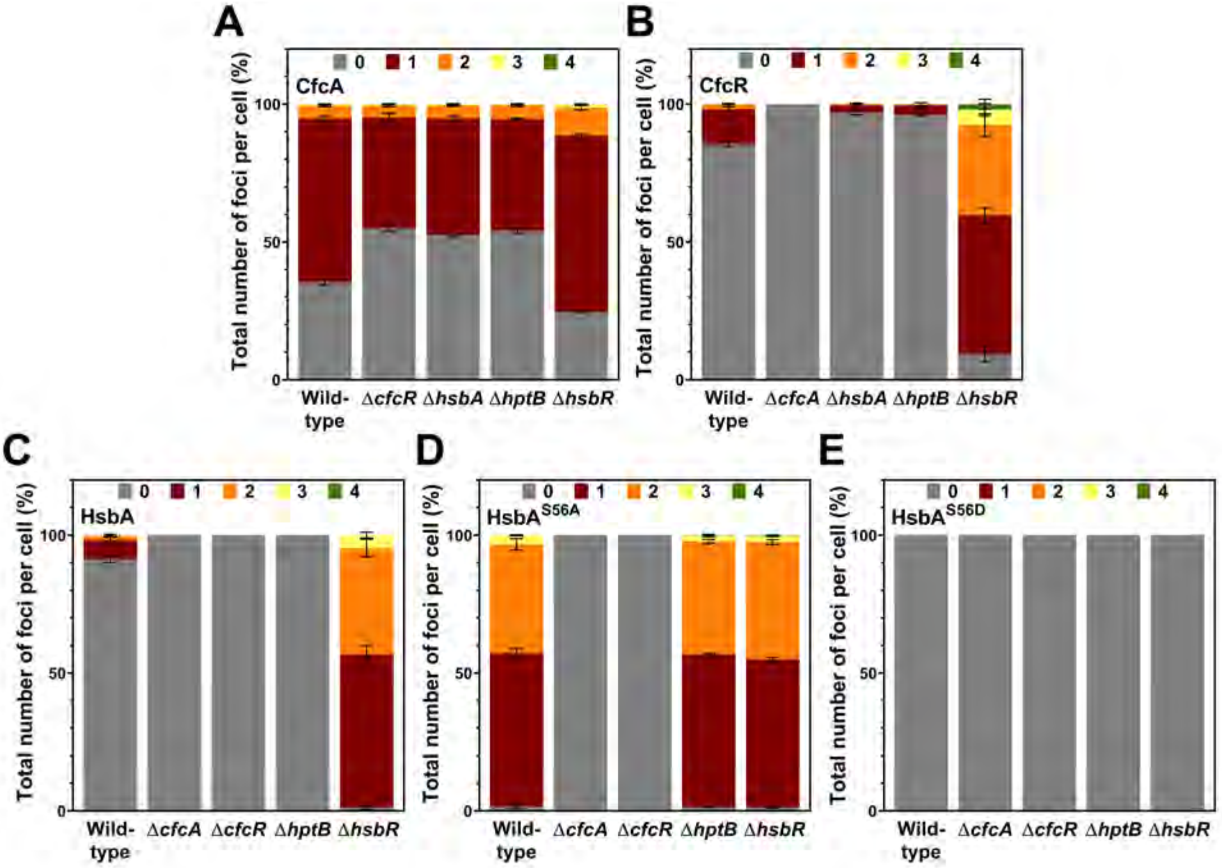


**Figure S10. Quantitative analysis of CfcA, CfcR and HsbA foci.** Percentage of wild-type, Δ*cfcR,* Δ*cfcA,* Δ*hsbA,* Δ*hptB* or Δ*hsbR* cells bearing 0, 1, 2, 3 or 4 total CfcA-GFP **(A)**, CfcR-GFP **(B)**, HsbA-GFP **(C)**, HsbA$^{S56A}$-GFP **(D)** or HsbA$^{S56D}$-GFP **(E)** fluorescent foci. Column and error bars represent averages and standard deviations of at least three separate replicates (n=500 cells).

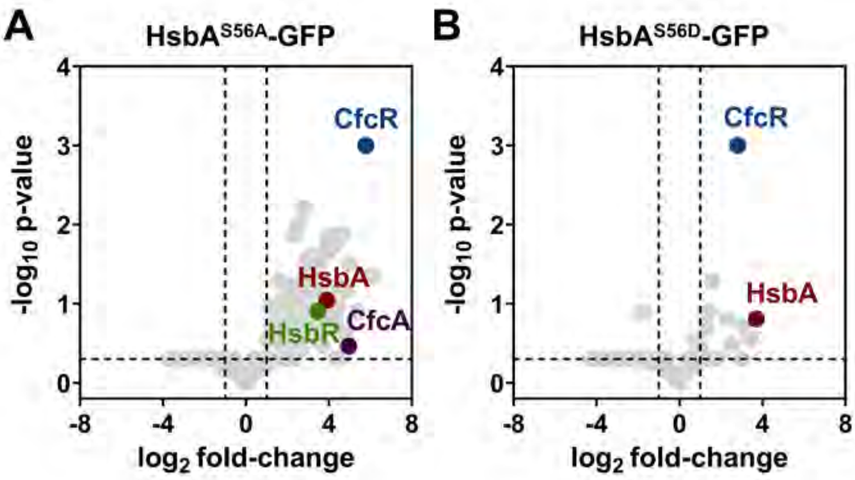

**Figure S11. Interactors of non-phosphorylatable and phosphomimetic HsbA.** Volcano plot showing the proteins co-immunoprecipitated with HsbA$^{S56A}$-GFP **(A)** or HsbA$^{S56D}$-GFP **(B)** in stationary-phase wild-type cells bearing the P*hsbA*$^{S56A}$-*hsbA-gfp* **(A)** or P*hsbA*$^{S56D}$-*hsbA-gfp* **(B)** transposons, respectively, as identified by mass spectrometry. Points show the $\log_2$ of the average fold-change in the peptide counts for each identified protein compared the negative control *vs.* the -$\log_{10}$ p-value of these peptide counts (n=2). Coloured dots indicate proteins assessed in this work.

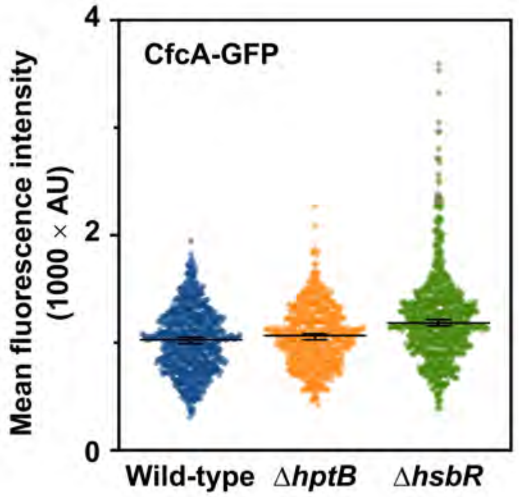


**Figure S12. Quantification of the CfcA-GFP fluorescent foci in the Δ*hptB* and Δ*hsbR* mutants.** Mean fluorescence intensity of CfcA-GFP foci in wild-type, Δ*hptB* and Δ*hsbR* cells (n=500 cells). Points correspond to individual fluorescence intensity values, bars denote the median and the 95% confidence interval of the median.

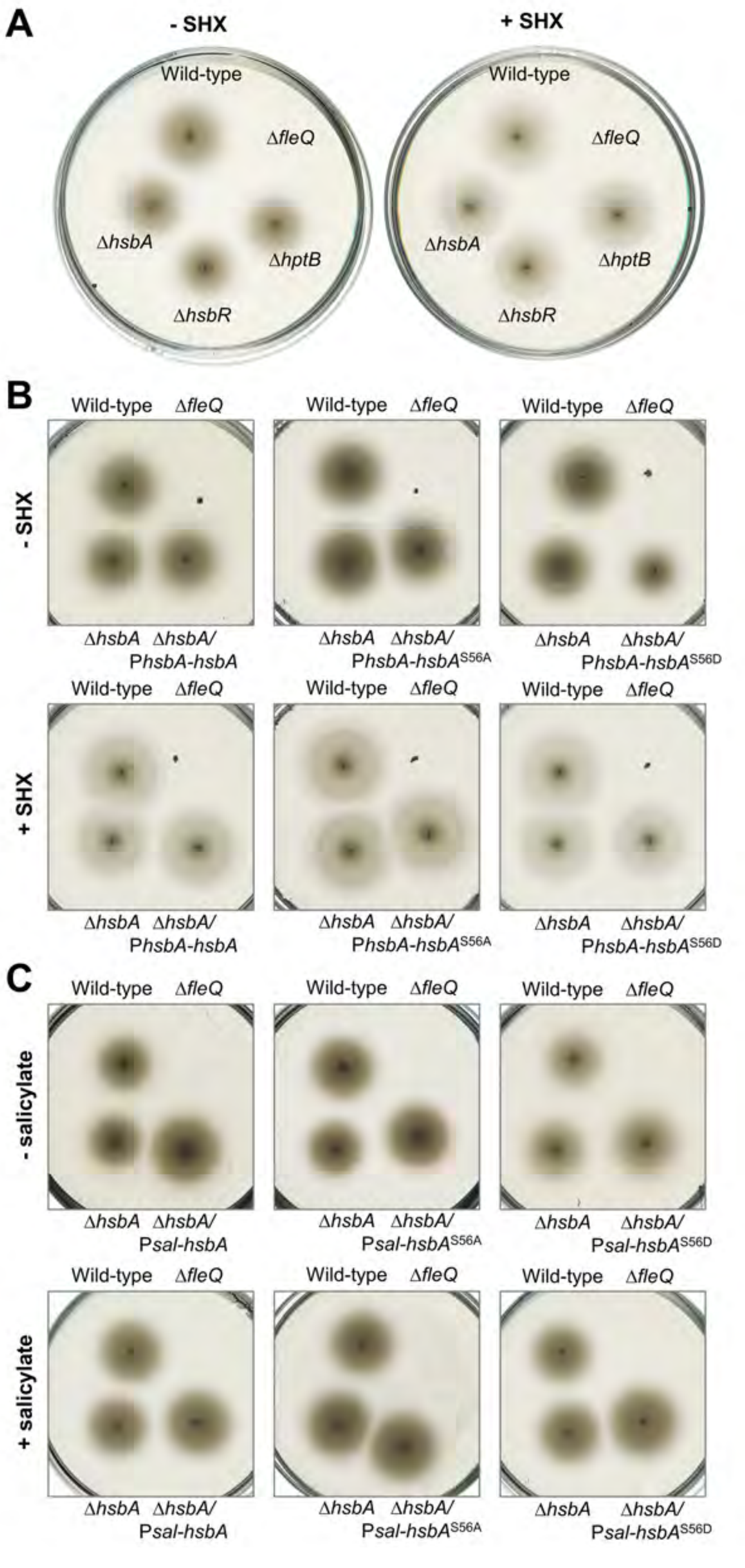


**Figure S13. Soft agar-based swimming motility halos.** Images of the swimming halos displayed by **(A)** the Δ*hsbA*, Δ*hsbR* and Δ*hptB* mutants in the absence (left) or the presence (right) of 0.8 mM SHX, **(B)** the Δ*hsbA* mutants bearing the P*hsbA-hsbA*, P*hsbA-hsbA*$^{S56A}$ or P*hsbA-hsbA*$^{S56D}$ transposons in the absence (top) or the presence (bottom) of 0.8 mM SHX, and **(C)** the Δ*hsbA* mutant bearing the P*sal-hsbA*, P*sal-hsbA*$^{S56A}$ or P*sal-hsbA*$^{S56D}$ transposons in the absence (top) or the presence (bottom) of 2 mM salicylate. The wild-type strain and Δ*fleQ* mutant were assayed in each plate as positive and negative controls, respectively. Each picture shows a representative swim plate out of at least three biological replicates.

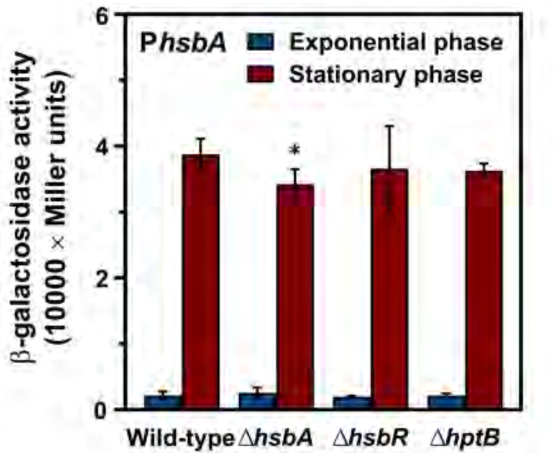


**Figure S14. Analysis of P*hsbA* expression.** β-galactosidase assays of exponential and stationary phase cultures of wild-type (KT2442), Δ*hsbA*, Δ*hsbR* and Δ*hptB* cells bearing a transcriptional P*hsbA*-*lacZ* fusion. Columns and error bars represent the averages and standard deviations of at least three biological replicates. Stars designate p-values for the Student's t-test for unpaired samples not assuming equal variance. ns: non-significant; *:$p<0.05$; **:$p<0.01$; ***:$p<0.001$; ****:$p<0.0001$.

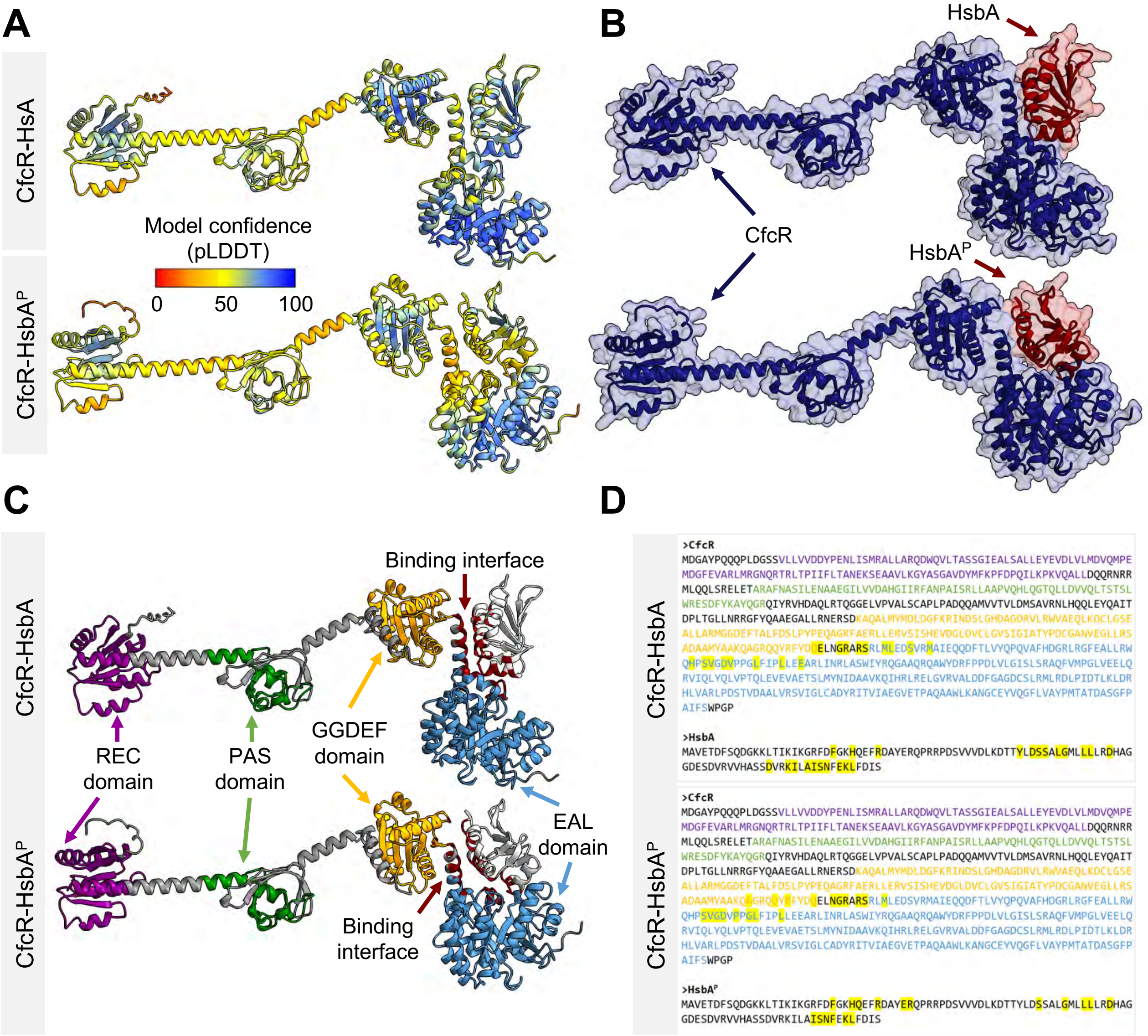


**Figure S15. Models of CfcR-HsbA and CfcR-HsbA-P complexes.** AlphaFold3-multimer prediction of the CfcR-HsbA and CfcR-HsbA-P interaction. Amino acid residues in cartoon and surface structures are colored to show **(A)** their predicted local distance difference test (pLDDT) score, representing the confidence of the predicted structure, **(B)** the protein they belong to (dark blue: CfcR; red: HsbA), and **(C)** locations of the CfcR REC (purple), PAS (green), GGDEF (yellow) and EAL (blue) domains, and the HsbA and CfcR residues located at the predicted binding interface (red). **D.** Amino acid sequences of CfcR and HsbA (top) or HsbA-P (bottom). Residues located at the binding interface are highlighted in yellow. Letter coloring as follows: REC domain (purple); PAS domain (green); GGDEF domain (yellow); EAL domain (blue).
